# PDZ and LIM domain protein 2 plays dual and context-dependent roles in breast cancer development

**DOI:** 10.1101/2020.01.27.920199

**Authors:** Josef Maryas, Jan Pribyl, Pavla Bouchalova, Petr Skladal, Pavel Bouchal

**Author notes:** Corresponding author: Pavel Bouchal, Ph.D. Masaryk University Faculty of Science, Department of Biochemistry Kamenice 5, 62500 Brno Czech Republic.

## Abstract

**Background:** PDZ and LIM domain protein 2 (PDLIM2) is a cytoskeletal and nuclear effector that regulates the activity of several transcription factors (e.g., NF-κB, STAT), and its deregulation has been associated with oncogenesis. Our recent study identified PDLIM2 as a protein associated with the lymph node metastasis of low grade luminal A breast cancer tissues. Here, we aim to understand this association at the molecular and cellular levels.

**Methods:** To investigate the link between PDLIM2 and epithelial-to-mesenchymal transition (EMT), stably transduced MCF7-PDLIM2 cells, and MCF7 or MCF10A cells with PDLIM2 protein levels modified using siRNA or *PDLIM2* gene carrying plasmid, were used. Additionally, MCF7 and MCF10A cells were exposed to hypoxic conditions and TGFβ1 treatment. EMT was monitored using immunoblotting of EMT markers and atomic force microscopy (AFM). The role of PDLIM2 in cell migration and/or invasion was investigated using Transwell assay and xCELLigence system.

**Results:** First, we observe a positive effect of PDLIM2 overexpression on EMT in MCF7 cells, a model of luminal A tumors, using EMT markers and AFM. On the other hand, PDLIM2 helps to maintain the epithelial phenotype in MCF10A cells, a model of normal breast epithelial cells. Second, we find that exposure of the MCF7 cells to hypoxic conditions increases levels of PDLIM2 and carbonic anhydrase-9 (CA-9), a marker of the response to hypoxia. However, none of these effects are observed in the MCF10A cells. Third, PDLIM2 overexpression promotes migration, invasion, and proliferation and decreases adhesion of the MCF7 cells, but an opposite effect is observed in the MCF10A cells.

**Conclusions:** Our data indicate that PDLIM2 plays a dual role: (i) as an EMT-supporting and hypoxia-responding oncoprotein in luminal breast cancer cells, and (ii) as an epithelial phenotype-maintaining tumor suppressor in normal epithelial breast cells.

## BACKGROUNDS

Breast cancer is the most common form of cancer and the second cause of death of women worldwide. Distant metastases represent the main reason for patient mortality [1, 2]. For risk group discrimination and determination of the metastatic potential of breast tumors in clinical practice, both traditional and molecular prognostic markers have been used. However, currently available markers are not sufficient for precise determination of metastatic potential [3, 4]. This insufficiency is well demonstrated by the low grade luminal A breast cancer subtype, whose general prognosis is favorable; nevertheless, approximately one-third of these tumors exhibit early lymph node metastases in disagreement with the initial prognosis. The molecular mechanism responsible for this lapse is unknown [3]. To improve understanding, we identified a panel of proteins associated with lymph node metastasis in low grade luminal A breast cancer using combined proteomics and transcriptomics [5]. PDZ and LIM domain protein 2 (PDLIM2), also known as Mystique or SLIM, was one of the key proteins in this panel, and its mRNA and protein levels were upregulated in a set of 24 lymph node-positive luminal A grade 1 tumors compared to their 24 lymph node-negative counterparts, specifically in this subtype. Additionally, the PDLIM2 protein level was higher in grade 1 vs. grade 3 tumors and in HER2^-^ vs. HER2^+^ breast cancer in a set of 96 breast tumors in total, indicating its connection with the luminal A subtype, low tumor grade, and high metastatic potential.

PDLIM2 is a member of the actinin-associated LIM protein (ALP) family, which contains a single N-terminal PDZ domain and C-terminal LIM domains [6]. All members of the ALP family are known to interact with the actin cytoskeleton [7] and play essential roles in its organization, cell differentiation, organ development, and neural signalization and have been associated with oncogenesis in general [8]. PDLIM2 acts as an E3 ubiquitin ligase and as such regulates the stability of NF-κB and other transcription factors in hematopoietic and epithelial cells. Its deregulation has been associated with several malignancies [7, 9–10]. The *PDLIM2* gene is localized on chromosome 8p21, a region frequently disrupted in various cancers [10–11], and its expression has been previously connected with both tumor suppression and oncogenesis [10]. *PDLIM2* expression is epigenetically suppressed by promoter hypermethylation in different cancers [10], and its re-expression is able to inhibit tumorigenicity and induce tumor cell death both *in vitro* and *in vivo* [12]. It was first identified in corneal epithelial cells [6] and in fibroblasts transformed by overexpression of the insulin-like growth factor 1 receptor (IGF-1R), T-lymphocytes, macrophages, dendritic cells and epithelial cancer cells [6-7, 9-11]. On the other hand, *PDLIM2* is highly expressed in invasive cancer cells, and its expression is associated with tumor progression and metastasis [5, 9].

Controversial literature results on the role of PDLIM2 in cancer development led us to assumptions about its dual and context-dependent role [13]. To test this hypothesis, we performed a series of *in vitro* experiments at the molecular and cellular levels using MCF7 breast cancer cells and MCF10A immortalized normal epithelial breast cells. Comparison of migration and invasion abilities as well as association with epithelial-to-mesenchymal transition (EMT) and hypoxia of different cell lines with modulated levels of PDLIM2 confirmed our hypothesis regarding the dual and context-dependent roles of PDLIM2 in breast cancer. Based on our data, we postulated a precise hypothesis about the cyclic changes of PDLIM2 protein levels during the oncogenesis of breast cancer, which also explains the inconsistency among previously published studies.

## MATERIALS AND METHODS

### Reagents and antibodies

Pyruvate, bromophenol blue, glycerol, mercaptoethanol, acrylamide, TEMED, APS, Tween 20, SDS, Tris HCl, HEPES, PEI, TGF β1, luminol, coumaric acid, resazurin, insulin, CH_3_COONa, NaBO_3_.4H_2_O, NaCl, KCl, Na_2_HPO_4_.12H_2_O, KH_2_PO_4_, MgCl_2_, and NaH_2_PO_4_ were purchased from Sigma-Aldrich. Streptomycin/penicillin, trypsin and EGTA were purchased from Invitrogen. Fetal bovine serum and horse serum were purchased from Biochrom AG, GoldView^TM^ was obtained from Viswagen, EDTA from Serva, EGF from Millipore, Calcein AM from Biotium, crystal violet from Merck, hydrocortisone from VUAB Pharma, and formaldehyde from Penta. Mouse anti-E-cadherin (diluted 1:100) antibody (Ab) and anti-vimentin Ab (1:1000) were purchased from DakoCytomation, mouse anti-p65 Ab (1:500) was purchased from Santa Cruz Biotechnology, mouse anti-N-cadherin Ab was purchased from Invitrogen, mouse anti-actin Ab (1:250) was purchased from Sigma Aldrich and mouse anti-PDLIM2 Ab (1:250) was purchased from Origene. Rabbit anti-FAK Abs (1:500), anti-β-catenin Ab (1:500) and anti-p-p53 (phosphorylated on S20) Ab (1:500) were purchased from Cell Signaling. Mouse anti-CA9 Ab (M75, diluted 1:3), anti-KRT18 Ab (DC10, 1:10), anti-p53 (DO-1, 1:10) and anti-PCNA Ab (PC10, 1:10) were prepared in-house. Horseradish peroxidase-conjugated RAMPx and SWARPx Abs (both DakoCytomation, dilution 1:1000) were used as secondary Abs. All antibodies were diluted in PBS with 0.1% Tween 20 containing 5% nonfat milk.

### Cell lines, cell culture and cell counting

The human ER-positive breast cancer cell line MCF7 and ER-negative breast cancer cell lines MDA-MB-231 and BT-549 were maintained in DMEM supplemented with 10% FBS, 1.25 mM pyruvate, 0.172 mM streptomycin and 100 U/ml penicillin. The human nontumorigenic breast epithelial cell line MCF10A was cultivated in DMEM/F12 supplemented with 5% horse serum, 1.25 mM pyruvate, 0.172 mM streptomycin, 100 U/ml penicillin, 20 ng/ml EGF, 0.5 mg/ml hydrocortisone and 10 µg/ml insulin. All cell lines were purchased from the American Type Culture Collection (ATCC). The cells were counted using a CASY TT cell counting device (Roche, Mannheim, Germany) or Bürker chamber for adhesion, proliferation, migration and invasion experiments.

### Commercial plasmids and siRNAs

For specific plasmid preparation, the pENTR221 “entry” vector and pcDNA3-GW-DEST “destination” vector were used (both from Invitrogen). Empty pcDNA3-GW-DEST served also as a CTRL plasmid in the experiments. On-Target plus SMART pool human PDLIM2 siRNA was used for PDLIM2 suppression (600 nM concentration, cat. no. L-005152-00). As a control, CTRL siRNA On-Target plus Nontargeting pool control siRNA (cat. no. D-001810-10-20) was used also at a concentration of 600 nM (both siRNAs from Dharmacon, Thermo Scientific).

### Plasmid construction

#### Total RNA isolation and One step RT PCR

Total RNA was isolated from the BT-549, MDA-MB-231 and MCF7 cell lines using an RNeasy Mini Kit 250 (Qiagen) according to the manufacturer’s protocol. The isolated RNA was pooled, eluted into 50 µl of RNase-free water, and stored at -80 °C. Complementary DNA (cDNA) was synthesized and amplified using a One Step RT PCR kit (Invitrogen) with PDLIM2-specific primers: forward 5’-CGGCGCCGGGCTCCTCTC-3‘ and reverse 5’-CCCTGGCCCACCCCTCTCCTTCC-3’. The reaction mixture contained 1 M betain, 1x Reaction Mix, 0.36 M trehalose, 1 µl of SuperScript^TM^ III RT/Platinum Taq Hi Fi Enzyme Mix, 0.5 µg of total RNA, and primers at a concentration 0.4 µM, and nuclease-free water was added to the final volume of 50 µl. The PCR program included two steps of cDNA synthesis (50 °C for 30 min and 80 °C for 3 min), predenaturation at 95 °C for 30 s, and 10 amplification cycles - each consisting of denaturation at 95 °C for 5 s, annealing at 55 °C for 30 s and extension at 68 °C for 3 min, followed by further extension at 68 °C for 1 min. The resulting product was purified using the standard protocol of the QIAquick PCR Purification Kit (Qiagen).

#### Construction of the pcDNA3-PDLIM2-GW-DEST plasmid

Gateway technology (Thermo Fisher Scientific) was used to construct the PDLIM2-carrying plasmid. The cDNA fragment of PDLIM2 was amplified using PCR with Herculase II DNA polymerase, a pair of specific primers, PDLIM2 TEV forward (5’-GGCTCTGAGAACCTGTACTTCCAGAGCATGGCGTTGACGGTGGATGTG-3’, bearing a region coding the sequence recognized by TEV protease) and PDLIM2 GWs reverse (5’-GTACAAGAAAGCTGGGTTTCAGGCCCGAGAGCTGAGG-3’, bearing a stop codon), and a pair of universal primers, TEV forward (TEV F: 5’-GGGGCTGCTTTTTTGTACAAACTTGTCCGAGACTCTTGG-3’) and ATTB2 reverse (ATTB2 R: 5’-GGGGCAGCTTTCTTGTACAAAGTGGGACATGTTCTTTCG-3’). The reaction mixturel of Herculase II, specific contained 1 M betain, 20µl of cDNA, 1x reaction buffer, 1 mM dNTP, 1 μM of Herculase II, specific primers at 0.1 µM and universal primers at 0.3 µM, and nuclease-free water was added to the final volume of 50 µl. The PCR program included predenaturation at 95 °C for 1 min and 30 amplification cycles - each consisting of denaturation at 95 °C for 10 s, annealing at 50 °C for 20 s and extension at 70 °C for 90 s, followed by further extension at 70 °C for 5 min. The final product was separated by electrophoresis on a 1.5% GoldView^TM^ stained agarose gel and visualized under UV light by a CCD camera. The target fragment was extracted by a QIAquick Gel Extraction Kit (Qiagen) according to the manufacturer’s protocol, and 150 ng were used for the BP recombinant reaction with pDONR221 according to the Gateway technology manual (Thermo Fisher Scientific). Chemically competent *E. coli* TOP10 (Life Technologies) cells were used for the preparation and amplification of resulting entry vector (pENTR221-PDLIM2). The QIAprep Spin Miniprep Kit (Qiagen) was used for vector purification according to the manufacturer’s protocol. A total of 150 ng of purified and sequencing-verified entry vector together with 150 ng of pcDNA3-GW-DEST vector were used for preparation of the pcDNA3-PDLIM2-GW-DEST destination plasmid, and the LR recombination reaction was performed according to the Gateway technology manual (Thermo Fisher Scientific). Chemically competent *E. coli* TOP10 cells were used again for preparation and amplification of the resulting destination plasmid that was subsequently purified by the Qiagen Plasmid Maxi Kit (Qiagen). The resulting destination plasmid was stored at -20 °C.

### Construction of lentiviral plasmids and lentiviruses and generation of a stably transduced MCF7-PDLIM2 cell line

The lentiviral vector pLENTI-PDLIM2 was prepared in-house according to Gateway® Technology with the Clonase® II user guide (Invitrogen, 25-0749 MAN0000470). The production of lentiviruses was performed according to the ViraPower™ Lentiviral Expression Systems user manual (Invitrogen, 25-0501 MAN0000273). Transduction of MCF-7 cells and selection of stably transfected clones were performed according to the ViraPower™ Lentiviral Expression Systems user manual (Invitrogen, 25-0501 MAN0000273).

### Cell transfection

For PDLIM2 suppression, cells were transfected using the Amaxa cell line nucleofector kit V (LonzaBio). A total of 1.0×10^6^ cells cultivated to 70% confluence were harvested and resuspended in 100 µl of AMAXA buffer (4 mM KCl, 10 mM MgCl_2_, 120 mM NaH_2_PO_4_/Na_2_HPO_4_, 10 mM HEPES pH 7.2) together with either anti-PDLIM2 siRNA or control siRNA at a concentration of 600 nM. T024 and P020 transfection programs were used for the transfection of MCF 10A and MCF7 cells, respectively. Plasmid transfection using the polyethylene imine (PEI) method based on lipofection was used. Cells were cultivated on 6-cm Petri dishes to 60-70% confluence, and 5 µg of specific or control plasmid was resuspended together with 15 µl of PEI (working solution 1 µg/µl in water, pH 7) in 0.5 ml of serum-free medium, incubated for 15 minutes at room temperature, poured onto a dish and further cultivated for 24, 48 or 72 hours.

### SDS PAGE and immunoblotting

Cell lysates for SDS PAGE were prepared using hot (95 °C) sample buffer (10% glycerol, 2% bromophenol blue, 62.5 mM Tris HCl pH 6.8, 2% SDS pH 6.8, 5% mercaptoethanol). SDS PAGE with a 5% stacking gel and 10% running gel was used for separation as described previously [14]. Protein lysates in the amount of 20 µ g, determined by an RC-DC Protein Assay (Bio-Rad), were run in the gels and wet-transferred onto PVDF membranes. Membranes were then blocked for 1 hour in PBS+0.1% Tween 20 (2.68 mM KCl, 0.137 M NaCl, 6.45 mM Na_2_HPO_4_.12H_2_O, 1.47 mM KH_2_PO_4_,

0.89 mM Tween 20) containing 5% nonfat milk and incubated with primary antibody (at the appropriate dilution, see above) at 4 °C overnight. After incubation, membranes were washed two times in PBS and once in PBS+0.1% Tween 20 and subsequently incubated with the corresponding secondary antibody (1:1000) at room temperature for 1 hour and washed again. After 5 min of incubation of membranes with ECL solution (10 mM luminol, 0.5 mM EDTA, 405 µM coumaric acid, 200 mM Tris pH 9.4, 8 mM sodium perborate tetrahydrate, 50 mM sodium acetate), immunoreactive proteins were visualized by enhanced chemiluminescence (ECL) solution using a CCD camera (Alpha Innotech FluorChem^TM^SP, Quansys Biosciences, USA).

### Young’s modulus mapping by Atomic Force Microscopy

Young’s modulus mapping was performed using JPK NanoWizard 3 (JPK, Berlin, Germany) on a bioAFM microscope, similar to previous publications [15–17]. A non-coated silicon nitride AFM probe Hydra 2R 100N (AppNano, Mountain View, CA, USA) equipped with a pyramidal silicon tip was used for all experiments (side angle 18°). The sensitivity and cantilever spring constant were calibrated by the recommended routine procedure. The 64x64 point maps of force distance curves (setpoint 1 nN, Z length 15 µ m, time per curve 0.5 s) were measured. The recorded FD curves were fitted with the Bilodeau modification [18] of the Hertzian model in AtomicJ software [19]. Final visualization of the images and stiffness maps was performed with Gwyddion software ver. 2.44 [20]. Further details are provided in the Additional file 15: Methods S1.

### TGFβ1 treatment and hypoxia

EMT in cells was induced by TGFβ1 treatment or hypoxic conditions. For PDLIM2 protein level induction and monitoring, TGFβ1 was added to the complete culture medium to a final concentration of 1 ng/ml, and the cells were cultivated for 24 hours. For cell morphology changes and monitoring, TGF β1 at 10 ng/ml was added to serum-free medium after 16 hours of serum starvation, and the cells were cultivated for 48 hours [21]. Control cells were cultivated in medium without TGFβ1. For induction of EMT by hypoxia, cells were cultivated in a hypoxic culture hood (Biospherix Xvivo X3, Biospherix) under a 2% concentration of O_2_ for 48, 72 or 96 hours.

### Migration and invasion assay

Real-time measurements of cell migration and invasion were performed using the CIM-Plate 16 module of an xCELLigence System RTCA DP real-time cell analyzer (Roche, UK). For invasion measurement, the top side of the polycarbonate membrane in the upper chamber wells was coated with Culturex®Basement membrane extract (Trevigen, USA) diluted 1:50 with Coating buffer (Trevigen, USA) 4 hours prior to the experiment. The CIM plates were then prepared by the addition of 170 µL of corresponding medium (with 10% FBS as the chemoattractant) into twelve wells of the lower chamber; four wells filled with 170 µ L of serum-free medium served as controls to determine the background signal. Each well of the upper chamber was filled with 30 µ L of serum-free medium. The plates were inserted into the xCELLigence station in the culture hood (21% O_2_, 5% CO_2_, 37 °C), and the baseline impedance was measured after 1 hour of equilibration at 37 °C. Cells (5×10^4^ cells per well for migration and 1×10^5^ cells per well for invasion) were then added to the wells of the upper chamber in 100 µL of serum-free medium, and the plate was equilibrated again for 30 min at room temperature and inserted into the xCELLigence once again before the start of measurements. The detailed arrangement of the plates is shown in Additional file 1: Figure S1. The xCELLigence analyzer was set to measure the impedance every 15 min for 24 hours. Normalized cell indexes (CI) were then calculated as a variable corresponding to the number of migrating or invading cells and statistically evaluated (part “Statistical analysis”).

Migration and invasion of cells were also measured using a 96-well Transwell assay (Trevigene, USA). The Transwell assay consists of upper and lower chambers (each with 96 wells) separated by a porous polyethylene terephthalate (PET) membrane (pore size 8μm) localized on the bottom of the upper chamber wells. For migration measurement, 100 µl of medium containing 10% FBS (used as a chemoattractant) was applied to the lower chamber wells, and wells were filled with 100 μl of serum-free medium were used as controls. Cells (5×10^4^ cells per well for migration and 1×10^5^ cells per well for invasion; see Additional file 1: Figure S1 for plate design) were resuspended in serum-free medium and added to the upper chamber wells. After incubation in a culture hood (21% O_2_, 5% CO_2_, 37 °C) for 24 hours, upper and lower chamber wells were rinsed, cells in the upper chamber wells were washed away and the migrated or invasive cells on the lower membrane surface were stained using Calcein AM according to the manufacturer’s protocol. After 1 hour of incubation in the culture hood in the dark, the fluorescence of converted calcein was measured using a Tecan Infinite M100 Pro (Life Sciences) with an excitation wavelength of 495 nm and emission wavelength of 515 nm. The number of cells migrating or invading across the membrane was quantified according to fluorescence values and statistically evaluated (part “Statistical analysis”). The invasion assay was performed equivalently to the migration measurement assay but with coating of the top side of the polycarbonate filter in the upper chamber wells with Culturex® Basement membrane extract.

Each experiment was performed with two biological replicates (cells grown on independent plates) per condition with three technical replicates (wells) for each biological replicate. Two independent experiments were performed per comparison.

### Adhesion assay

To measure adhesion in monolayer culture, cells in corresponding medium were seeded at 1×10^5^ cells per well in triplicate in a 6-well plate. After 30 min, the medium was completely removed from the wells, and unattached cells in the medium were counted using a Bürker chamber. The adhesion of cells was based on the amount of unattached cells, which was statistically evaluated (part “Statistical analysis”). Each experiment was performed three times on independent plates per comparison.

### Proliferation assay

To measure proliferation in monolayer culture, cells in corresponding medium were seeded at 2×10^4^ cells per well in triplicate in a 96-well plate. After 24 hours, cells were treated with resazurin (1 mg/ml) and incubated for 150 min. After the incubation period, the fluorescence of resorufin was measured using a Tecan Infinite M100 Pro, and measurement was performed for 16 spots per well (excitation wavelength 573 nm/emission wavelength 583 nm) and statistically evaluated. Each experiment was performed three times on independent plates per comparison.

### Statistical analysis

STATISTICA software version 12 was used for the statistical analyses for Figs. 3A-B and 5A. Data were reported as the mean +/-1.96*(standard deviation) corresponding to 95% confidence intervals; Student’s t-test was used to assess the significance of differences between two groups, and p-values below 0.05 were considered significant. For AFM evaluation in Fig. 1E, the Mann-Whitney test was performed in R version 3.5.3., and p-values below 0.05 were considered significant. For the remaining statistical analyses, Student’s t-test in Microsoft Excel was applied, and p-values below 0.05 were considered significant.

**Fig. 1:**
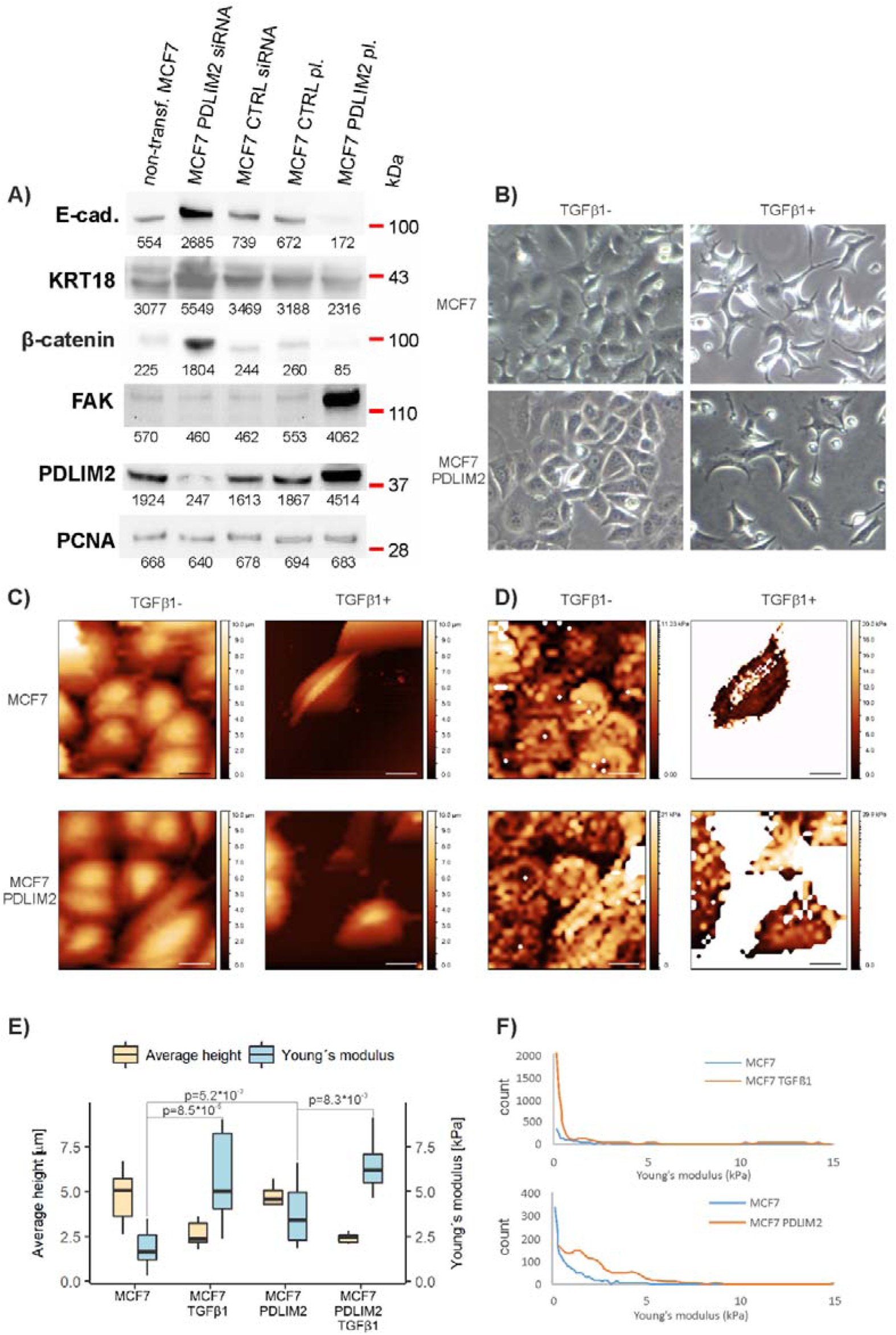
Effect of PDLIM2 protein level modulations on EMT markers and MCF7 cell stiffness. (A) Effect of modulated PDLIM2 levels on the EMT markers E-cadherin (E-cad.), keratin-18 (KRT18) and β-catenin as well as focal adhesion kinase (FAK) in MCF7 cells. PDLIM2 protein levels were modulated by siRNA suppression (MCF7 PDLIM2 siRNA) compared to the control (MCF7 CTRL siRNA) or by PDLIM2 overexpression (MCF7 PDLIM2 pl.) compared to the control (MCF7 CTRL pl.). Proliferating cell nuclear antigen (PCNA) was used as a loading control [25]. Numbers under the protein bands represent their integral optical density (INT*mm^2^). Blots are representative of two independent experiments (biological replicates), see Additional file 2: Figure S2 for both biological replicates. **(B)** Representative photos of MCF7 parental cells and stably transduced MCF7-PDLIM2 cells after TGFβ1 treatment (TGFβ1+) in comparison with control untreated cells (TGFβ1-). Magnification 50x. **(C) Di**stribution of cell height measured by atomic force microscopy (AFM) over the MCF7 parental cells and stably transduced MCF7-PDLIM2 cells after TGFβ1 treatment (TGFβ1+) in comparison with control untreated cells (TGFβ1-). Scale bar inside the im e is equal to 20 µm. Figures are representative of 11 AFM measurements (biological replicates) per group, see Additional file 16: Dataset S1 for all measurements (also applies to Fig. 1D). **(D)** Distribution of Young’s modulus measured by AFM over the MCF7 parental cells and stably transduced MCF7-PDLIM2 cells after TGFβ1 treatment (TGFβ1+) in comparison with control untreated cells (TGFβ1-). **E)** Average height and young’s modulus of the cells measured by AFM: MCF7 and MCF7-PDLIM2 cells after TGFβ1 treatment (TGFβ1+) in comparison with control untreated cells (TGFβ-) Statistics for the key Young’s modulus comparisons are shown, see Tab. S1 for full statistics. Eleven AFM measurements (biological replicates) per group. **F)** Histograms show distribution of Young’s modulus in MCF7-PDLIM2 and TGFβ1-treated MCF7 cells compared to parental/untreated MCF7 cells.

## RESULTS

### PDLIM2 protein levels are positively coregulated with epithelial-to-mesenchymal transition in MCF7 breast cancer cells

Based on the results of our previous combined proteomics-transcriptomics study [5], we hypothesized that PDLIM2 may act as an oncoprotein in luminal A breast tumors. To test this hypothesis *in vitro*, we selected MCF7 human breast cancer cells as a model of luminal A breast cancer [22–24], modulated PDLIM2 protein levels and evaluated molecular and cellular changes caused by these modifications. First, overexpression of PDLIM2 decreased the epithelial markers E-cadherin, β-catenin and keratin-18 (KRT18) and increased levels of the β1-integrin pathway regulator FAK (Fig. 1A and Additional file 2: Figure S2), indicating epithelial-to-mesenchymal transition (EMT) was connected with PDLIM2. On the other hand, *PDLIM2* expression suppression by small interference RNA (siRNA; Fig. 1A and Additional file 2: Figure S2) increased E-cadherin, β-catenin and KRT18 and decreased FAK levels, indicating mesenchymal-to-epithelial transition (MET) was initiated by PDLIM2 suppression. The ability of PDLIM2 to affect cell morphology was observed in MCF7 cells stably transduced with *PDLIM2* vector (compared to parental MCF7 as a control) using optical microscopy, showing that MCF7-PDLIM2 cells acquire similar morphology as MCF7 cell after TGFβ1-induced EMT (Fig. 1B). A similar pattern was evident from the atomic force microscopy (AFM) data, showing MCF7-PDLIM2 cells had higher stiffness based on Young’s modulus relative to parental MCF7 cells (Fig. 1E), similarly as after TGFβ1-induced EMT (Figs. 1B-F, Additional file 12: Table S1 and Additional file 16: Dataset S1). All of these data suggest that PDLIM2 might play a significant role in the regulation of EMT in MCF7 cells.

### EMT induction by TGFβ1 and hypoxia increases PDLIM2 levels, and PDLIM2 overexpression augments CA9 and reduces p53 levels as well as p53 phosphorylation (S20) in MCF7 breast cancer cells

To investigate how PDLIM2 is affected by EMT, we induced EMT by TGFβ1 (1 ng/ml for 24 hours) or by long-term exposure to hypoxia (2% O_2_ for 96 hours) and monitored how PDLIM2 responds. Both treatments led to elevated PDLIM2 protein levels, and successful induction of EMT was confirmed by decreased levels of the epithelial markers E-cadherin, β-catenin and KRT18 (Fig. 2A and Additional file 4: Figure S4). Notably, we observed that shorter exposure times to hypoxic conditions (48 and 72 hours) were still not able to decrease levels of epithelial markers and induce EMT (see Additional file 5: Figure S5); however, PDLIM2 levels increased (Fig. 2B and Additional file 5: Figure S5), indicating that not only EMT but also hypoxia itself upregulates PDLIM2 levels. Moreover, levels of carbonic anhydrase-9 (CA9), a marker of the response to hypoxic conditions, were elevated in MCF7 cells with overexpressed PDLIM2 (Fig. 2C and Additional file 6: Figure S6).

**Fig. 2:**
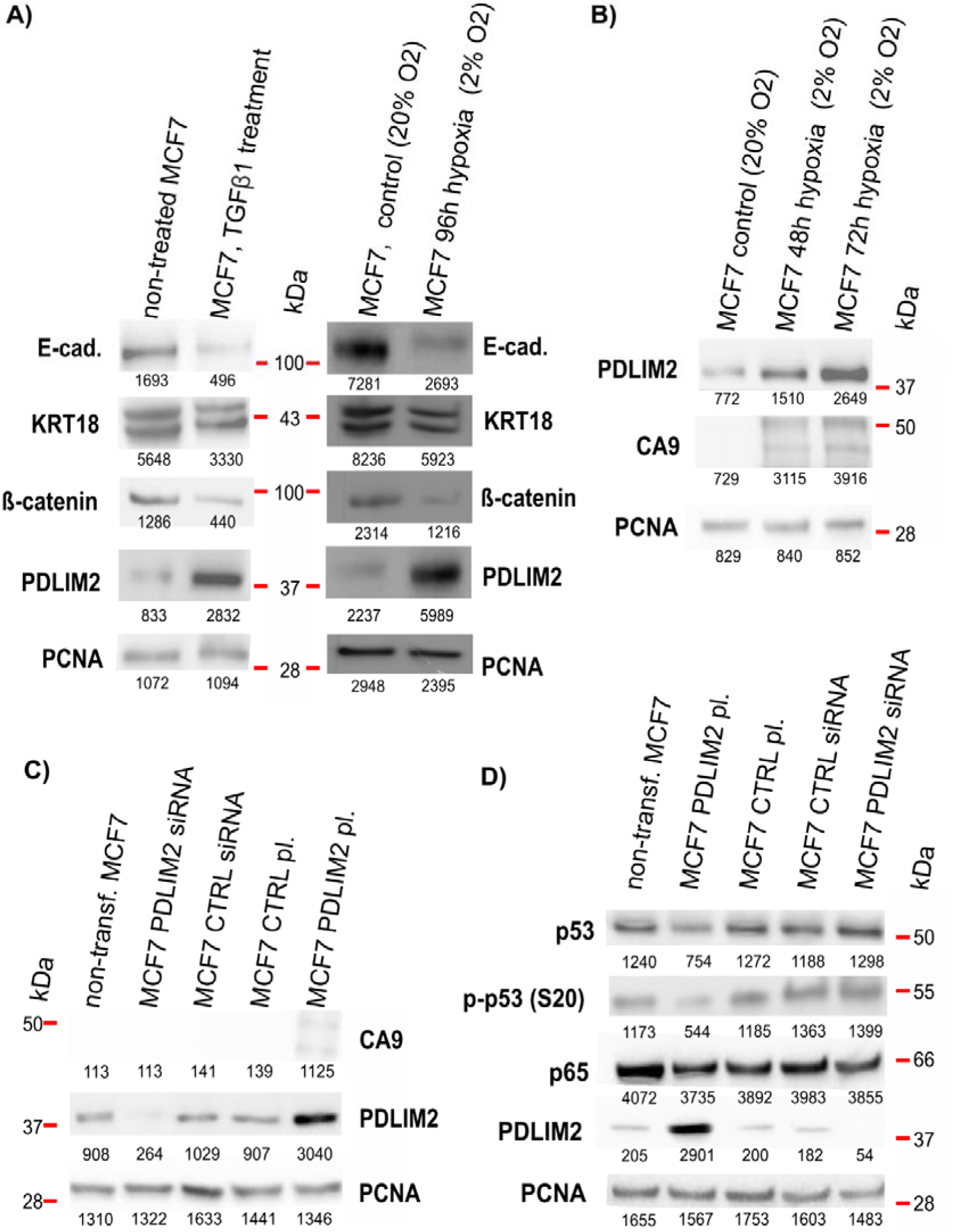
Effect of EMT on PDLIM2 levels and, conversely, PDLIM2 modulations on selected proteins (MCF7 cells). **(A)** Effect of EMT induction by TGFβ1 treatment (1 ng/ml for 24 hours) and by long-term hypoxia (96 h) on PDLIM2 protein levels. Successful EMT induction was confirmed by changes in the EMT markers E-cadherin (E-cad.), keratin-18 (KRT18) and β-catenin. **(B)** Effect of short-term hypoxia (48 and 72 hours) on PDLIM2 protein levels. Carbonic anhydrase-9 (CA9) was monitored as a control marker of hypoxia induction. **(C)** Effect of PDLIM2 overexpression on CA9 protein levels. **(D)** Effect of PDLIM2 protein level modulations on p53 protein levels, p53 (S20) phosphorylation (p-p53 S20), and on p65. Proliferating cell nuclear antigen (PCNA) was used as a loading control. Numbers under the protein bands represent their integral optical density (INT*mm^2^). Blots are representative of two independent experiments (biological replicates), see Additional files 4 - 7: Figures S4 - S7 for both biological replicates.

In addition to TGFβ1 and hypoxia-induced EMT, we were interested in whether PDLIM2 functionally interacts with other key cancer players, including p53 and p65. We found that PDLIM2 overexpression decreased both the p53 total protein level as well as its serine 20 phosphorylated, active form (p-p53 (S20)) (Fig. 2D and Additional file 7: Figure S7). On the other hand, the level of p65 protein, a key member of the canonical NF-κB pathway, was not affected by any of these treatments, nor by overexpression or suppression of PDLIM2 (see Fig. 2D and Additional file 7: Figure S7).

### PDLIM2 overexpression increases both migration and invasion of MCF7 breast cancer cells

To further verify the pro-tumorigenic role of PDLIM2 on cellular levels, we examined its effect on the migration and invasion of the MCF7 cells. PDLIM2 overexpression increased the migration capabilities of these cells as observed by real-time measurement using the xCELLigence system (p=2.8x10^-4^) and independently confirmed using a Transwell assay with end-point detection (p=3.6x10^-5^) (Fig. 3A). Overexpression of PDLIM2 also had a similar effect on the invasion of MCF7 cells measured using xCELLigence (p=3.9x10^-4^) and the Transwell assay (p=1x10^-5^) (Fig. 3B). Our results suggest that PDLIM2 is relevant for regulation of the migration and invasiveness of MCF7 cells.

**Fig. 3:**
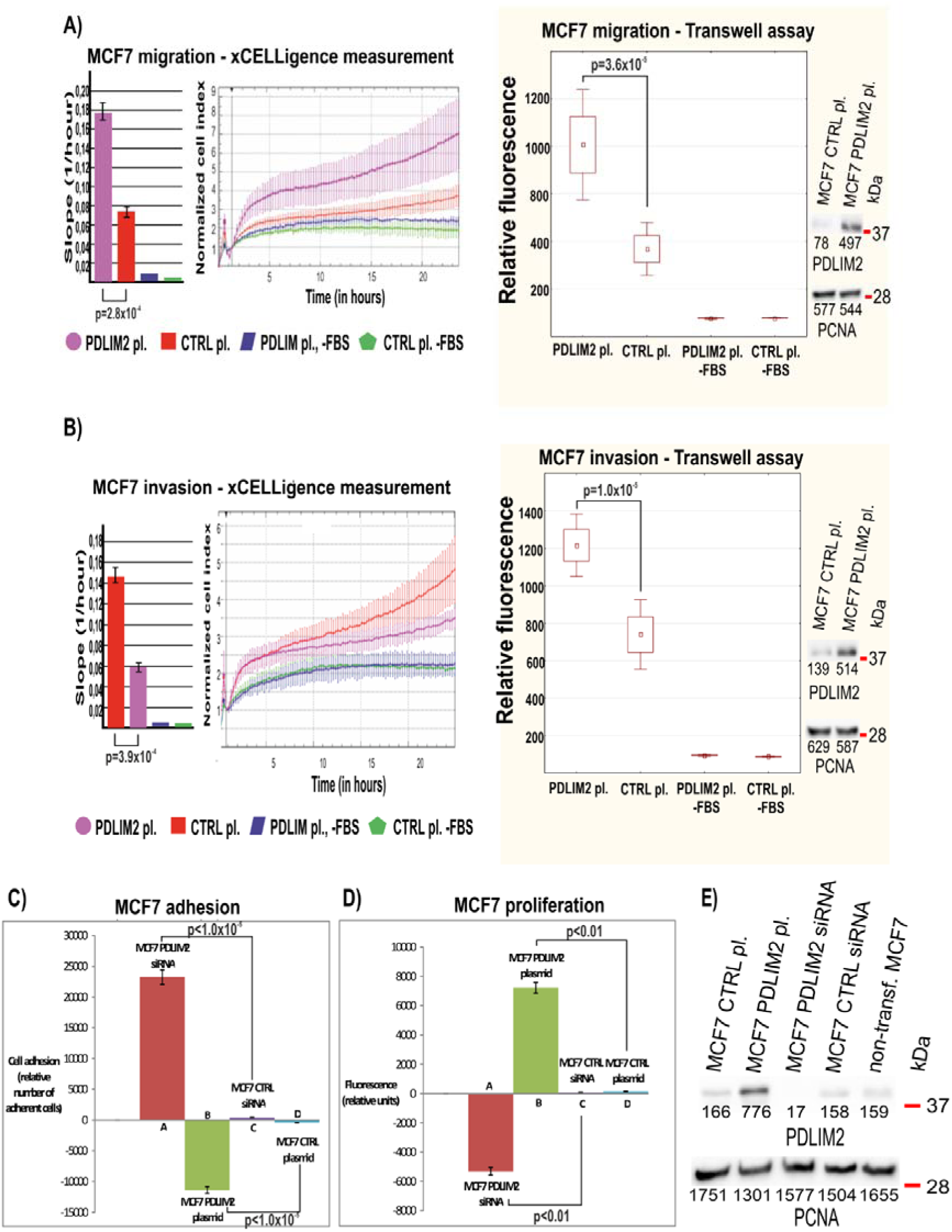
Effect of PDLIM2 protein level modulations on migration, invasion, adhesion and proliferation of MCF7 cells. **(A)** Effect of PDLIM2 overexpression (PDLIM2 pl.) on the migration of MCF7 cells (in comparison with control cells with endogenous PDLIM2 levels (CTRL pl.)) measured by the xCELLigence system and Transwell assay. Conditions without fetal bovine serum as the chemoattractant (-FBS) serve as negative controls in A and B. **(B)** Effect of PDLIM2 overexpression (PDLIM2 pl.) on the invasiveness of MCF7 cells measured by the xCELLigence system and by Transwell assay. **(C)** Effect of PDLIM2 protein level modulations on the adhesion of MCF7 cells. PDLIM2 protein levels were modulated by siRNA suppression (MCF7 PDLIM2 siRNA) compared to the control (MCF7 CTRL siRNA) or by PDLIM2 overexpression (MCF7 PDLIM2 pl.) compared to the control (MCF7 CTRL pl.) **(D)** Effect of PDLIM2 protein level modulations on the proliferation (viability) of MCF7 cells. **(E)** Verification of PDLIM2 protein level modulations in MCF7 cells measured in C and D; please see separate verification blots for independently cultivated cells in A and B figure sections/measurements. PCNA was used as a loading control. Numbers under the protein bands represent their integral optical density (INT*mm^2^). All data were obtained from at least two independent experiments (see Additional files 17 - 18: Datasets S2 - S3 for migration and invasion experiments). For the detailed design of the xCELLigence and Transwell assay plates see Additional file 1: Figure S1.

### PDLIM2 overexpression diminishes cell adhesion and increases proliferation of MCF7 cells, while suppression of PDLIM2 has the opposite effect

Next, we investigated the consequences of PDLIM2 modulation on the adhesion and proliferation of MCF7 cells. Suppression of PDLIM2 by siRNA significantly augmented the adhesion of MCF7 cells (*p=1x10^-5^) compared to control cells; on the other hand, specific overexpression of PDLIM2 considerably decreased adhesion (p=1x10^-5^) (Fig. 3C). PDLIM2 suppression also substantially diminished the proliferation of MCF7 cells (p=1x10^-5^), in contrast to the dramatic increase in the proliferation of the MCF7 cells with overexpressed PDLIM2 (p=1x10^-5^) (Fig. 3D). These results indicate involvement of PDLIM2 in the regulation of MCF7 cell adhesion and proliferation.

### PDLIM2 overexpression is important for maintenance of the epithelial phenotype in MCF10A human immortalized epithelial breast cells, while PDLIM2 suppression has opposite consequences

To ascertain whether the effects related to PDLIM2 levels described in sections above are of general validity or are context-dependent, we selected another model, MCF10A immortalized human epithelial breast cells. We either overexpressed or suppressed PDLIM2 in these cells and monitored the effects of these changes on molecular and cellular levels. The effects of overexpression and suppression of PDLIM2 on EMT markers are shown in Fig. 4A and Additional file 8: Figure S8: overexpression led to augmentation of the epithelial marker E-cadherin and decreases in the mesenchymal markers vimentin and N-cadherin, as well as the β1-integrin pathway regulator FAK, indicating MET and maintenance of the epithelial phenotype. PDLIM2 suppression had the opposite effect: E-cadherin levels were decreased, and levels of the mesenchymal markers vimentin and N-cadherin as well as FAK were increased, indicating EMT. These results suggest that PDLIM2 is important for maintaining the epithelial phenotype of MCF10A cells.

**Fig. 4:**
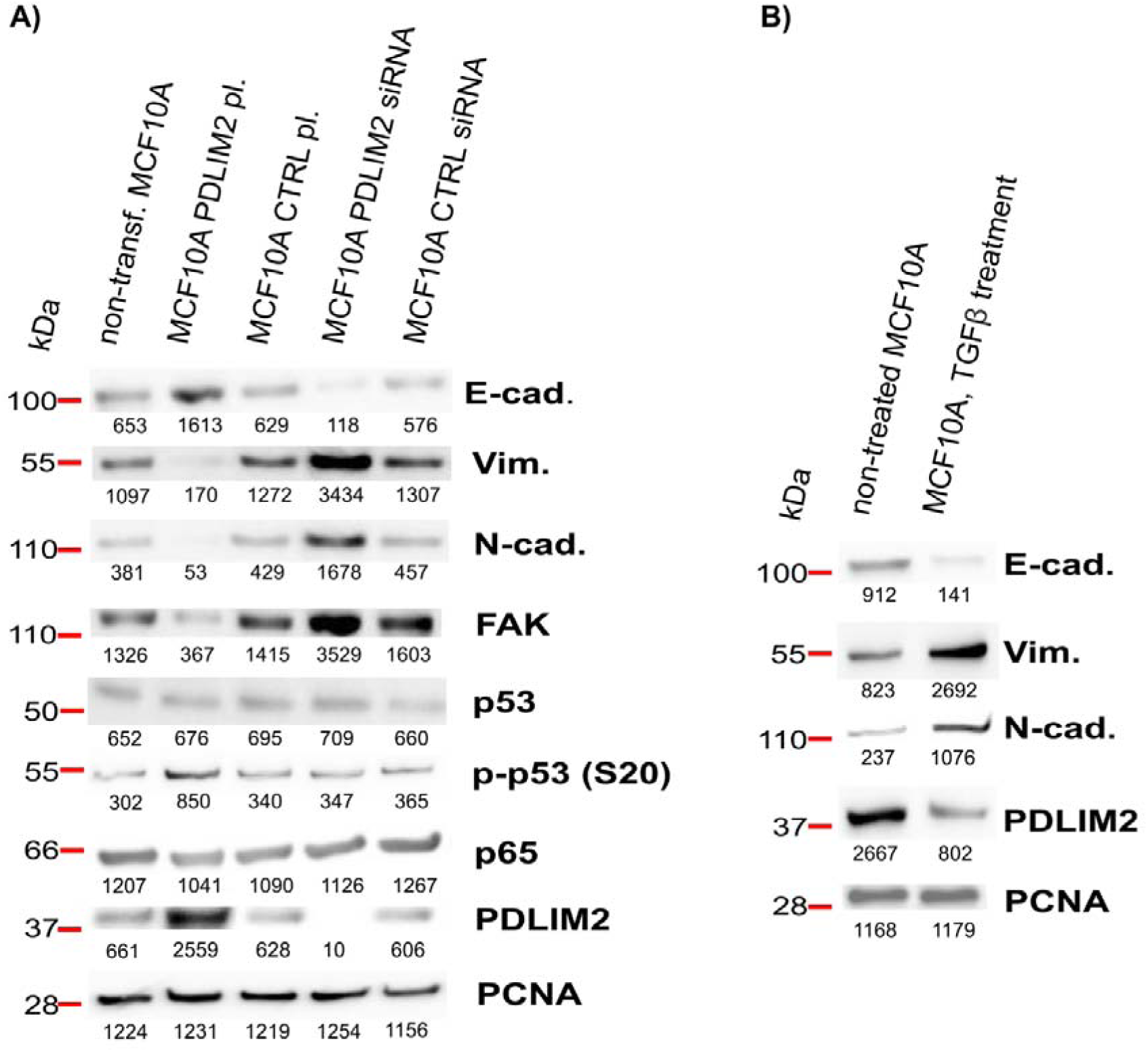
Effects of PDLIM2 modulations on EMT, and *vice versa* (MCF10A cells). **(A)** Effects of PDLIM2 protein level modulations on the EMT markers E-cadherin (E-cad.), N-cadherin (N-cad.) and vimentin (Vim.) as well as FAK levels, p53, p53 (S20) phosphorylation and p65. **(B)** Effects of EMT induction by treatment with TGFβl (1 ng/ml for 24 hours) on PDLIM2 protein levels and EMT markers. Numbers under the protein bands represent their integral optical density (INT*mm^2).^ Blots are representative of two independent experiments (biological replicates), see Additional files 8 - 9: Figures S8 - S9 for both biological replicates.

### EMT induction by TGFβ1 decreases PDLIM2 levels, but hypoxia has no effect on PDLIM2levels in MCF10A cells

To investigate how EMT affects PDLIM2 in MCF10A cells, we induced EMT by treatment with TGFβ1 (1 ng/ml for 24 hours) and monitored PDLIM2 protein levels. As shown in Fig. 4B and Additional file 9: Figure S9, PDLIM2 protein levels were downregulated after EMT induction in MCF10A cells, in a distinct manner compared to that of MCF7 cells. Successful EMT induction was confirmed by decreased levels of E-cadherin and increased levels of vimentin and N-cadherin. We also attempted to induce EMT by exposing the MCF10A cells to hypoxic conditions; however, we did not observe any changes in E-cadherin and vimentin levels as well as in PDLIM2 protein levels (see Additional file 10: Figure S10A and S10B). Additionally, we did not observe any change in CA9 levels after PDLIM2 modulation (see Additional file 10: Figure S10C). These results confirm the importance of PDLIM2 in the maintenance of the epithelial phenotype and indicate that PDLIM2 does not play a significant role in the response to hypoxic conditions in MCF10A cells.

### PDLIM2 overexpression decreases p53 (S20) phosphorylation in MCF10A cells

To investigate mechanisms of the potential tumor suppressive role of PDLIM2 in MCF10A cells, we analyzed its effect on p53 levels, p53 phosphorylation (S20) and p65 protein levels. In contrast to MCF7 cells, overexpression of PDLIM2 had no effect on p53 levels (see Fig. 4A and Additional file 8: Figure S8); nevertheless, the active form of this protein, p-p53 (S20), was augmented (Fig. 4A and Additional file 8: Figure S8). The level of p65 was not affected by overexpression or suppression of PDLIM2 (see Fig. 4A and Additional file 8: Figure S8). These results suggest a tumor suppressive role of PDLIM2 in MCF10A cells.

### PDLIM2 levels negatively regulate migration and positively regulate the adhesion of MCF10A cells

To validate the distinct role of PDLIM2 in MCF10A cells at the cellular level, we examined its effect on the migration and adhesion of MCF10A cells. PDLIM2 suppression by siRNA was accompanied by significant augmentation of the migration abilities of the MCF10A cells relative to cells with endogenous protein levels, as revealed by Transwell assay (p<1x10^-5^) (Fig. 5A, conditions A, C and E). Conversely, overexpression of PDLIM2 significantly decreased the migration abilities of the MCF10A cells (p<1x10^-5^) relative to cells with endogenous protein levels (Fig. 5A, conditions B, D and E). As shown in Fig. 5B, PDLIM2 suppression significantly decreased the adhesion of MCF10A cells (p<1x10^-5^), while on the other hand, overexpression of this protein considerably augmented the adhesion of MCF10A cells (p<1x10^-5^) relative to control cells. These results further confirm the tumor-suppressive role of PDLIM2 in MCF10A cells and its context-dependent behavior.

**Fig. 5:**
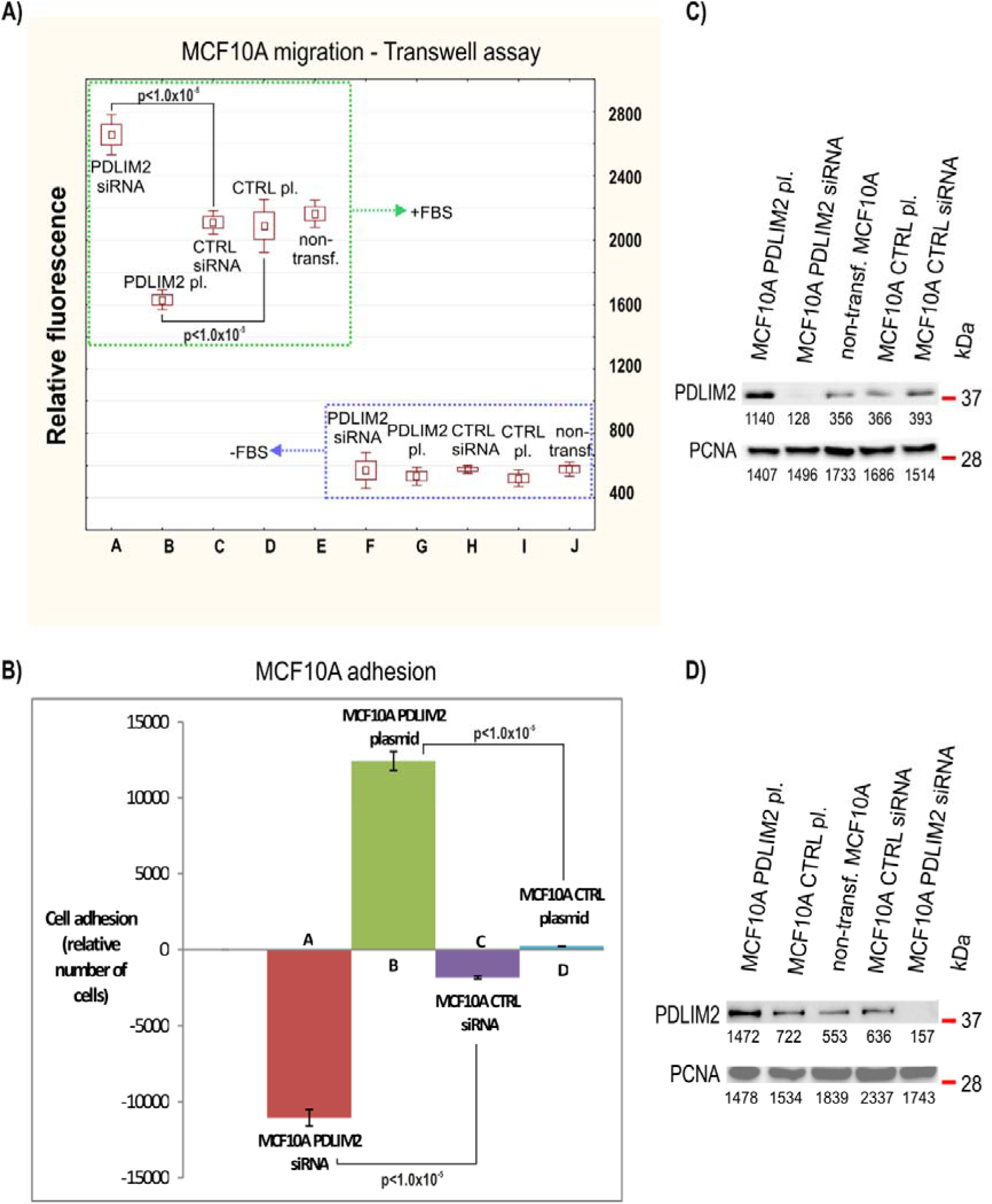
Effect of PDLIM2 protein level modulations on the migration and adhesion of MCF10A cells. **(A)** Effects of PDLIM2 protein level modulations on MCF10A cell migration. PDLIM2 protein levels were modulated by siRNA suppression (PDLIM2 siRNA) compared to the control (CTRL siRNA) or by PDLIM2 overexpression (PDLIM2 pl.) compared to the control (CTRL pl.) Conditions without fetal bovine serum as a chemoattractant (-FBS) serve as negative controls in A. **(B)** Effects of PDLIM2 protein levels on the adhesion of MCF10A cells. See legend to A for explanation. **(C)** and **(D)** Verification of PDLIM2 protein level modulations in MCF10A cells for A and B measurements, respectively. PCNA was used as a loading control. Numbers under the protein bands represent their integral optical density (INT*mm^2^). All data were obtained from at least two independent experiments (see Additional file 19: Dataset S4 for migration experiments).

### Analysis of breast cancer cell lines and tissues supports the dual role of PDLIM2 in breast cancer development

Finally, comparison of PDLIM2 protein levels in other breast cancer cell lines (Additional file 11: Figure S11) showed that PDLIM2 levels were low in low invasive MCF7 breast cancer; however, higher PDLIM2 levels were found in highly invasive triple negative MDA-MB-231, supporting the connection of PDLIM2 to metastatic potential. Higher levels of PDLIM2 were also found in normal MCF10A cells (Additional file 11: Figure S11), in agreement with its tumor suppressor role in normal breast cells. This is in principal agreement with our breast cancer tissue data that showed a high PDLIM2 level in primary tumors forming lymph node metastases, but specifically in small (T1) luminal A grade 1 tumors (Additional file 13: Table S2), which represent the early phase of tumor development. Conversely, the epithelial marker KRT18 was significantly downregulated in PDLIM2-rich luminal A tumors forming lymph node metastases (Additional file 14: Table S3), further supporting the PDLIM2 connection with mesenchymal phenotype in tissues. Taken together, these data suggest a connection between PDLIM2 and metastatic phenotype in luminal A tumors and its distinct roles in different phases of cancer development.

## DISCUSSION

Deregulation of PDLIM2 has been associated with oncogenesis, including lymph node metastasis of breast cancer [5]. The *PDLIM2* gene is repressed in different cancers, which implies its tumor suppressive role. Repression and a tumor suppressive role of PDLIM2 were observed in ATL induced by HTLV1 virus [26–29], Kaposi sarcoma [30], ovarian cancer [31], gastric cancer [32], colon cancer [12], Hodgkin lymphoma [33] and breast cancer [34, 35]. In contrast to the above studies, a pro-oncogenic role and enhanced expression of PDLIM2 were observed in castration-resistant prostate cancer cells [5, 9, 36], invasive breast cancer cell lines and breast carcinomas [9, 11]. The aim of the study presented here was to understand the role of PDLIM2 at both the molecular and cellular levels in the breast cancer context in more detail using *in vitro* experiments.

### PDLIM2 role in MCF7 breast cancer cells

Our data indicate a positive role for PDLIM2 in EMT induction (Fig. 1A and Additional file 2: Figure S2), which was previously described in DU145 prostate cancer cells and MDA-MB-231 highly invasive breast cancer cells [9, 36]. We for the first time report this role in MCF7 cells, a model of luminal A tumors, confirming the previous data from our clinical set of luminal A breast cancer tissues [5]. Overexpression of PDLIM2 also denotes disturbance of the β1-integrin pathway and loss of the epithelial phenotype *via* increased levels of FAK, the regulator of the β1-integrin pathway [10]. Apart from the above, the stiffness of MCF7 cells was significantly augmented after PDLIM2 overexpression (Fig. 1B-C). Increased cell stiffness is typical for cells with a mesenchymal phenotype [28], which again supports the PDLIM2 role in EMT induction. Moreover, EMT induction by TGFβ1 treatment and/or long-term hypoxia increases PDLIM2 levels (Fig. 2A, Additional file 4: Figure S4). Our experiments also revealed a reciprocal connection between PDLIM2 and the response to hypoxia (Fig, 2B, 2C, Additional file 5: Figure S5 and Additional file 6: Figure S6), which represents completely new information that deserves further examination. Furthermore, the observed negative effects of PDLIM2 on the levels of proteins involved in DNA repair and cell cycle regulation via p53 and especially p-p53 (S20) [39–41] indicate the relation between PDLIM2 and these processes in MCF7 cells (Fig. 2D and Additional file 7: Figure S7). All of these results confirm the involvement of PDLIM2 in the regulation of EMT induction and in the maintenance of the cellular phenotype in MCF7 cells. Additionally, involvement of PDLIM2 in the response to hypoxia and the effect of PDLIM2 overexpression on key cancer molecular players suggest a pro-oncogenic role of PDLIM2 in MCF7 cells, further validating our previous data from clinical tissues [5].

At the cellular level, we have clearly proven the positive effect of PDLIM2 on MCF7 cell migration, invasiveness, and proliferation and the negative effect on MCF7 adhesive abilities (Fig. 3A-D). Similar results were previously obtained with MCF7 [11], DU145 and MDA-MB-231 cells for migration [9] and with CRPC cells for both migration and invasion [36]. A negative effect of PDLIM2 on adhesion and a positive effect on proliferation were already described in DU145 and MDA-MB-231 cells [9], and the positive effect of PDLIM2 on proliferation was observed in CRPC cells [36]. Migration and invasion are closely connected to EMT induction, and both are the key steps in oncogenesis and metastasis formation [42–44]. The results observed at the cellular level are thus in good agreement with the molecular-level data. Additionally, disruption of cell adhesion and increased cell proliferation are the key signatures of oncogenesis [42]. In view of these facts, and in agreement with previous studies, the cellular-level results bring additional support for the pro-oncogenic role of PDLIM2 in luminal A breast cancer and functionally further validate our previous data from clinical tissues [5].

### PDLIM2 role in MCF10A normal epithelial breast cells

On the other hand, the potential tumor suppressive role of PDLIM2 was revealed in our experiments using immortalized normal epithelial MCF10A cells. Our data indicate a negative role of PDLIM2 in EMT induction, its importance for maintaining the epithelial phenotype of MCF10A cells and the negative effect of EMT on PDLIM2 protein levels (Figs. 4A, 4B, Additional file 8: Figure S8 and Additional file 9: Figure S9), distinct from MCF7 cells. These new findings are in good agreement with the known role of PDLIM2 in maintaining breast epithelial cell polarity [10]. Overexpression of PDLIM2 in MCF10A was further associated with increased p53 (S20) phosphorylation. This was revealed for the first time and indicates a positive connection between PDLIM2, DNA repair and cell cycle regulation in these cells (Fig. 4A, Additional file 8: Figure S8). Interestingly, no effect of PDLIM2 on hypoxia and no effect of hypoxia on PDLIM2 levels were observed, suggesting different and context-dependent roles of PDLIM2 in the response to hypoxic conditions (see Additional file 10: Figure S10). It seems that PDLIM2 helps to induce a hypoxic response and supports the proliferation of MCF7 cells; however, it may not be involved in hypoxic response regulation in MCF10A cells. In a broader context, PDLIM2 function in MCF10A cells is evidently highly distinct from that in MCF7 cells, being rather tumor suppressing. Furthermore, similar levels of PDLIM2 protein in invasive triple negative breast cancer cells (MDA-MB-231) and normal breast epithelial cells (MCF10A) (Additional file 11: Figure S11) indicate, in agreement with previous data [34], a significant role for PDLIM2 not only in MCF10A but also in invasive breast cancer cells. On the other hand, PDLIM2 levels were substantially lower in low invasive MCF7 cells (Additional file 11: Figure S11 and [34, 11]), suggesting that PDLIM2 is suppressed in a gap between neoplastically transformed and invasive breast cancer cells that undergo EMT. The tumor suppressive function of PDLIM2 in MCF10A cells was also evident in experiments at the cellular level. As shown for the first time here, PDLIM2 negatively regulates migration and positively regulates the adhesion of MCF10A cells (Fig. 5A-B). These effects are in agreement with the expected role of PDLIM2 in the maintenance of the epithelial phenotype and with its tumor-suppressive function in MCF10A cells. Altogether, these results give evidence for the dual and context-dependent role of PDLIM2, which acts as a possible tumor suppressor in normal epithelial cells (MCF10A) and as a potential oncoprotein in luminal A breast cancer (MCF7 cells), and suggest that PDLIM2 levels may fluctuate during oncogenesis.

### Context-dependent role of PDLIM2 in breast cancer progression

Data presented in this study led us to the hypothesis that changes in PDLIM2 protein levels during breast cancer oncogenesis are context-dependent (Fig. 6). According to our data and several previous studies [9-11, 36, 37], we assume that newly transformed low-invasion breast cancer cells have reduced levels of PDLIM2. Overexpression of PDLIM2 levels in these cells led to a dramatic increase in their metastatic potential and cancer progression, demonstrating the oncoprotein role of PDLIM2 (Figs. 1-3, Additional file 2: Figure S2, Additional file 4: Figure S4, Additional files 5-7: Figures S5-S7). According to our data (Additional file 11: Figure S11) and in agreement with others [9, 34], we further assume that highly invasive breast cancer cells express high levels of PDLIM2, and we expect that this could support them to form metastasis. However, these cells may undergo MET in the next step of the metastatic cascade, and PDLIM2 levels may diminish after successful metastatic colonization, supporting growth of the cells to build secondary tumors. In agreement with our previous tumor tissue data [5], luminal A grade 1 tumors with high levels of PDLIM2 represent a group of tumors with increased metastatic potential. Inhibition of PDLIM2 protein in these tumors has the potential to reduce migration, invasion and proliferation and augment adhesion properties due to phenotypic changes toward MET, which makes PDLIM2 an interesting target for further studies on its possible therapeutic application.

**Fig. 6:**
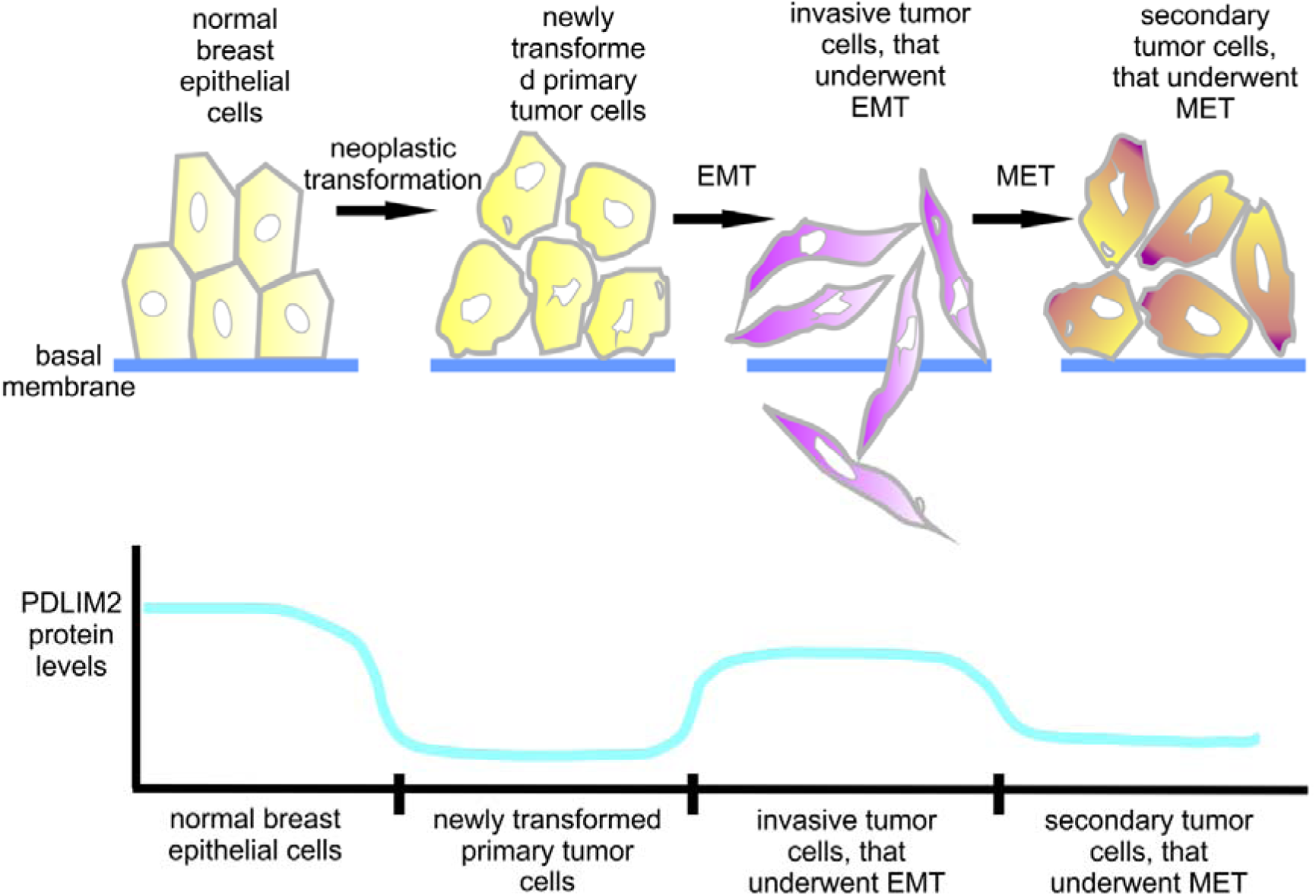
Model of context-dependent changes in PDLIM2 protein levels during breast oncogenesis. based on the data presented here. High PDLIM2 protein levels in normal epithelial breast cells are decreased during neoplastic transformation and are kept low in newly transformed tumor cells. However, PDLIM2 levels increase after EMT and are kept high in invasive tumor cells. Finally, PDLIM2 levels go down in secondary tumor cells that undergo MET and are kept as low as in newly transformed tumor cells.

## CONCLUSION

We demonstrate that PDLIM2 plays dual and context-dependent roles in breast cancer development: PDLIM2 facilitates maintenance of the epithelial phenotype in normal breast MCF10A cells, but it also promotes EMT in MCF7 breast cancer cells. PDLIM2 supports migration, invasion and proliferation in MCF7 cells but blocks migration and supports the adhesion of MCF10A cells. Our findings thus show that PDLIM2 has the potential to act as a tumor suppressor in MCF10A cells (a model of normal epithelial breast cancer) but as an oncoprotein in MCF7 cells (model of luminal A breast cancer). These observations are complementary and unravel previous contradictory findings in the literature, leading us to the hypothesis that changes in PDLIM2 protein levels during oncogenesis of breast cancer have a context-dependent nature. These data provide a basis for further interesting investigations to determine whether PDLIM2 blocking might have potential therapeutic implications in luminal A breast cancer.

## DECLARATIONS

### Ethics approval and consent to participate

Not applicable.

### Consent for publication

Not applicable.

### Availability of data and materials

All the datasets used and/or analysed during the current study are available from the corresponding author on reasonable request.

### Authors’ contributions

J.M. designed, performed and evaluated experiments and drafted the manuscript, J.P. performed and evaluated AFM experiments, P.B.^1^ prepared cells for optical microscopy and AFM and generated Fig. S3, P.S. initiated AFM experiments and contributed to manuscript preparation, P.B.^2^ supervised the study, manuscript preparation, acquired funding and approved the final manuscript.

## Acknowledgements

We thank Dr. Petr Müller for his help with preparation of the plasmids and stably transduced cell line, Zuzana Bertova for her technical assistance with AFM and Anna Pospisilova for data processing in Fig. 1E.

## ABBREVIATIONS

AFM: atomic force microscopy
ALP: actinin-associated LIM protein
CA9: carbonic anhydrase-9
E-cad: E-cadherin
EMT: epithelial-to-mesenchymal transition
FAK: focal adhesion kinase
IGF-1R: insulin-like growth factor 1 receptor
KRT18: keratin-18
MET: mesenchymal-to-epithelial transition
N-cad: N-cadherin
PCNA: proliferating cell nuclear antigen
PDLIM2: PDZ and LIM domain protein 2
Vim: Vímentin

## ADDITIONAL INFORMATION

**Additional file 1: Figure S1.pdf.** A common schema of xCELLigence and/or Transwell experiments for the measurement of cell migration and invasion.

**Additional file 2: Figure S2.pdf.** Effects of altered PDLIM2 protein levels on EMT markers in MCF7 cells.

**Additional file 3: Figure S3.pdf.** Confirmatory immunoblotting of EMT induction in MCF7-PDLIM2 and parental MCF7 cells after TGFβ1 treatment and before AFM measurements.

**Additional file 4: Figure S4.pdf.** Effect of EMT induction on PDLIM2 protein levels in MCF7 cells.

**Additional file 5: Figure S5.pdf.** Effects of short-term exposure to hypoxia (2% O_2_ for 48 and 72 hours) on EMT markers and PDLIM2 protein levels in MCF7 cells.

**Additional file 6: Figure S6.pdf.** Effects of altered PDLIM2 protein levels on the hypoxic marker carbonic anhydrase 9 (CA9) in MCF7 cells.

**Additional file 7: Figure S7.pdf.** Effects of altered PDLIM2 protein levels on p53 levels, p53 (S20) phosphorylation and p65 levels in MCF7 cells.

**Additional file 8: Figure S8.pdf.** Effect of altered PDLIM2 protein levels on EMT markers, p53, p-p53 (S20) and p65 in MCF10A cells.

**Additional file 9: Figure S9.pdf.** Effects of EMT induction on PDLIM2 protein levels in MCF10A cells.

**Additional file 10: Figure S10.pdf.** Effect of short term and long term expositions to hypoxic conditions on EMT induction and effect of PDLIM2 alterations on CA9 in MCF10A cells.

**Additional file 11: Figure S11.pdf.** PDLIM2 protein levels in different breast cell lines (MCF7, MCF10A and MDA MB 231).

**Additional file 12: Table S1.docx.** Statistics of average height and Young’s modulus of the cells measured by AFM.

**Additional file 13: Table S2.docx.** Connection between PDLIM2 and clinicopathological parameters of breast cancer based on our previous combined proteomics and transcriptomics study (n=96 in total) [5].

**Additional file 14: Table S3.docx.** iTRAQ-2DLC-MS/MS quantitative protein-level data for PDLIM2 and epithelial and mesenchymal marker proteins in lymph node-positive (n=24) vs. negative (n=24) luminal A grade 1 tumors in our previous combined proteomics and transcriptomics study (n=96 in total) [5].

**Additional file 15: Methods S1.pdf.** Young’s modulus mapping by Atomic Force Microscopy.

**Additional file 16: Dataset S1.pdf.** AFM data for all measurements.

**Additional file 17: Dataset S2.pdf.** xCELLigence data for MCF7 migration and invasion measurements.

**Additional file 18: Dataset S3.xls.** Transwell assay data for MCF7 migration and invasion measurements.

**Additional file 19: Dataset S4.xls.** Transwell assay data for MCF10A migration measurements.

