## Additional file 1 for "PDZ and LIM domain protein 2 plays dual and context-dependent roles in breast cancer development"

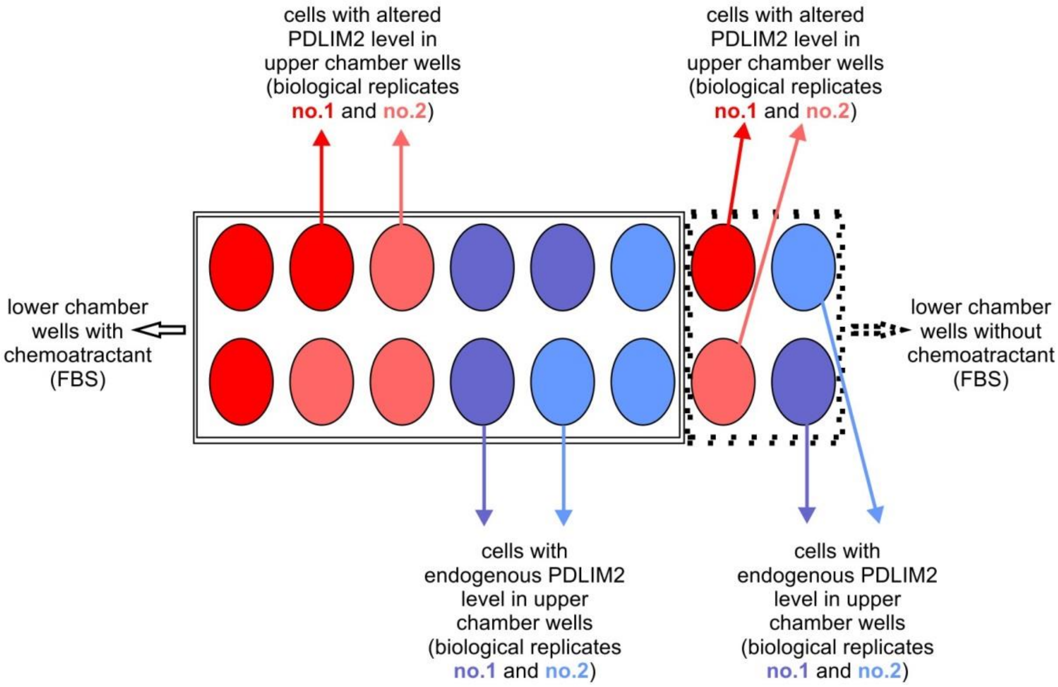


**Figure S1: A common schema of xCELLigence and/or Transwell experiments for the measurement of cell migration and invasion. Related to Figs. 3 and 5.** Eight wells in the upper chamber of the xCELLigence/Transwell desk contained cells with altered PDLIM2 levels (overexpression or downregulation; red), and another eight wells contained cells with endogenous PDLIM2 levels (blue). In both cases, four of eight wells contained cells grown as biological replicate no. 1 (dark), and another four of eight wells contained cells grown as biological replicate no. 2 (light; independent cultivations). Within each quartet of replicates, three wells were connected to lower chamber wells containing fetal bovine serum (FBS) as a chemoattractant (on the left), and one well was connected to the lower chamber without chemoattractant (on the right; negative control).
