## Additional file 2 for "PDZ and LIM domain protein 2 plays dual and context-dependent roles in breast cancer development"

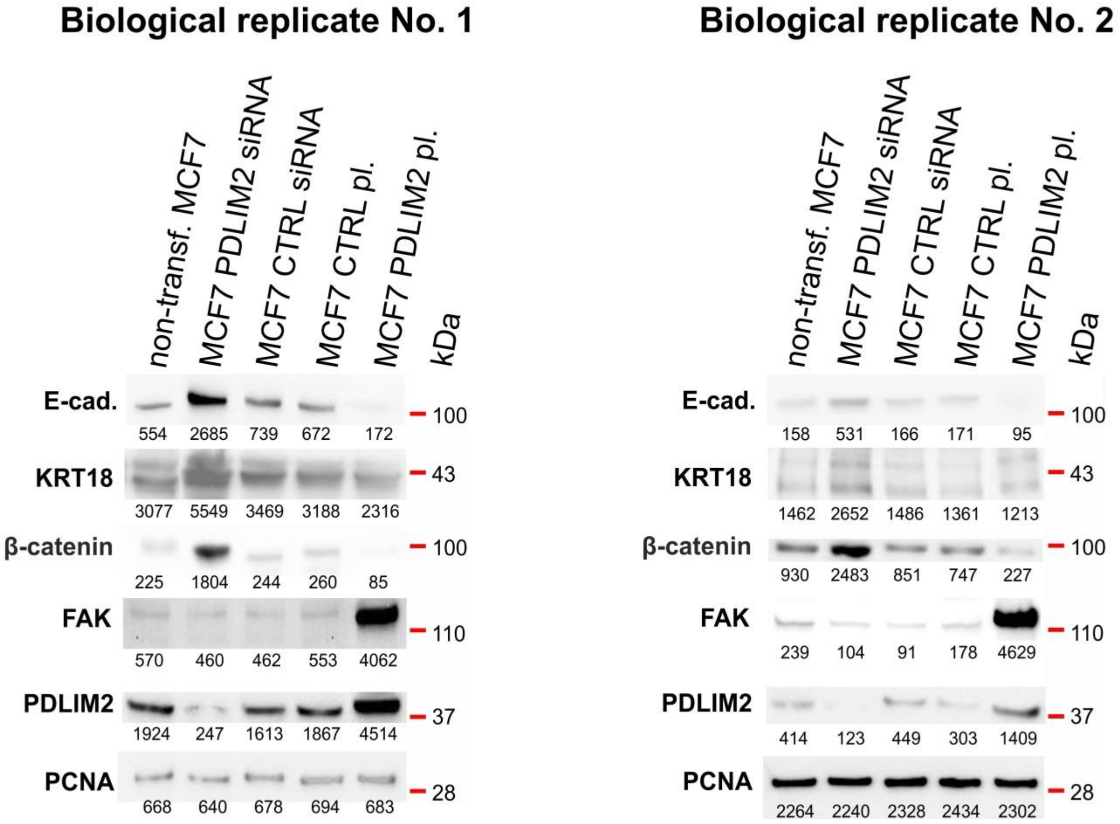


**Figure S2: Effects of altered PDLIM2 protein levels on EMT markers in MCF7 cells. Related to Fig. 1A.** PDLIM2 protein levels were altered by siRNA silencing (MCF7 PDLIM2 siRNA) compared to the control (MCF7 CTRL siRNA) or by PDLIM2 overexpression (MCF7 PDLIM2 pl.) compared to the control (MCF7 CTRL pl.). The EMT markers E-cadherin (E-cad.), keratin 18 (KRT18) and β-catenin as well as focal adhesion kinase (FAK) were monitored. PDLIM2 overexpression decreased the levels of the epithelial markers E-cadherin, KRT18 and β-catenin while increasing the levels of FAK; PDLIM2 silencing had the opposite effect. Blots represent two independent experiments (biological replicates). Numbers under the protein bands represent their integral optical density (INT*$mm^2$).
