## Additional file 3 for "PDZ and LIM domain protein 2 plays dual and context-dependent roles in breast cancer development"

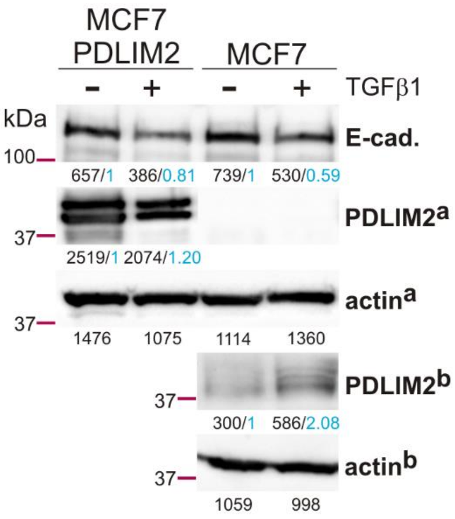


**Figure S3: Confirmatory immunoblotting of EMT induction in MCF7-PDLIM2 and parental MCF7 cells after TGFβ1 treatment and before AFM measurements. Related to Figs. 1B-F.** Levels of the epithelial marker E-cadherin (E-cad.) decreased, and PDLIM2 levels increased in both TGFβ1-treated MCF7-PDLIM2 and parental MCF7 cells in comparison with control cells. Black numbers under the protein bands represent their integral optical density (INT*mm$^2$), and blue numbers show actin normalized ratios in comparison with control cells. PDLIM2$^a$: 15 s exposure, PDLIM2$^b$, 3 min exposure.
