## Additional file 4 for "PDZ and LIM domain protein 2 plays dual and context-dependent roles in breast cancer development"

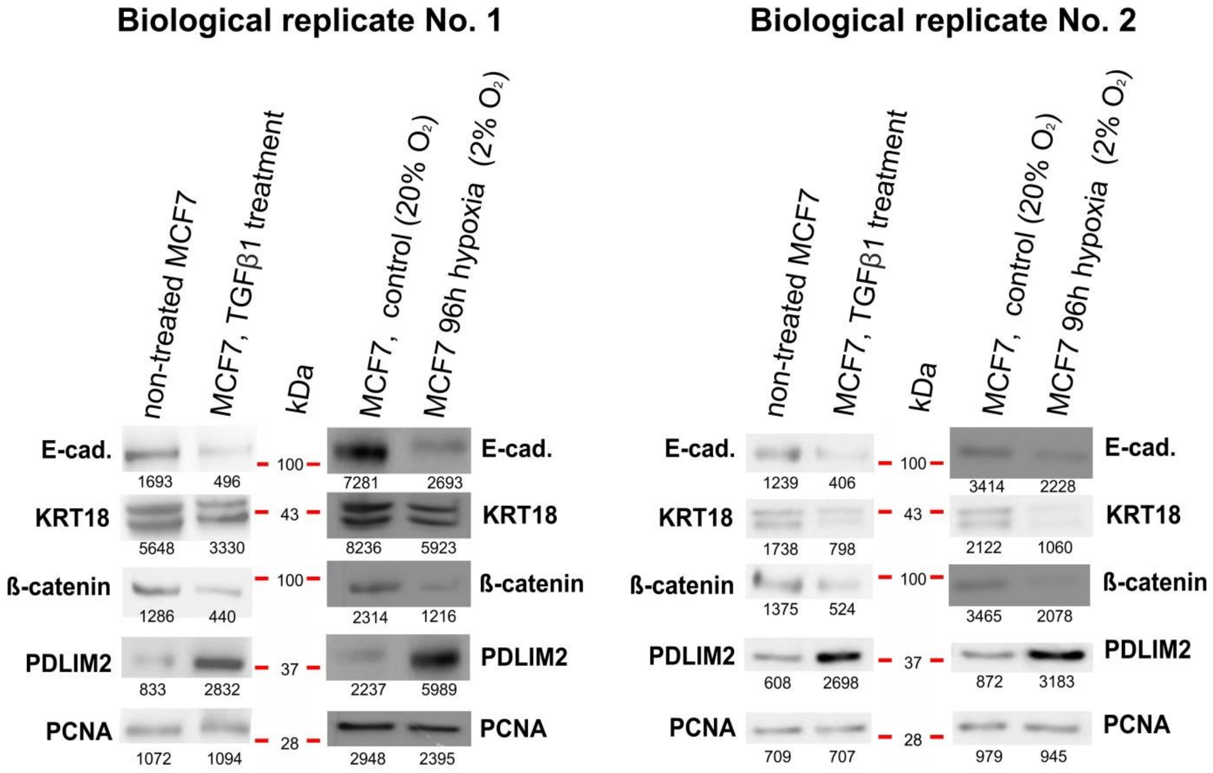


**Figure S4: Effect of EMT induction on PDLIM2 protein levels in MCF7 cells. Related to Fig. 2A.** EMT was induced either by TGFβ1 (1 ng/ml for 24 hours) or by long-term hypoxia (2% $O_2$ for 96 hours). EMT induction was monitored by the EMT markers E-cadherin (E-cad.), keratin-18 (KRT18) and β-catenin. Successful EMT induction led to increased levels of PDLIM2 and decreased levels of the epithelial markers E-cadherin, KRT18 and β-catenin. Blots represent two independent experiments (biological replicates). Numbers under the protein bands represent their integral optical density (INT*$mm^2$).
