## Additional file 5 for "PDZ and LIM domain protein 2 plays dual and context-dependent roles in breast cancer development"

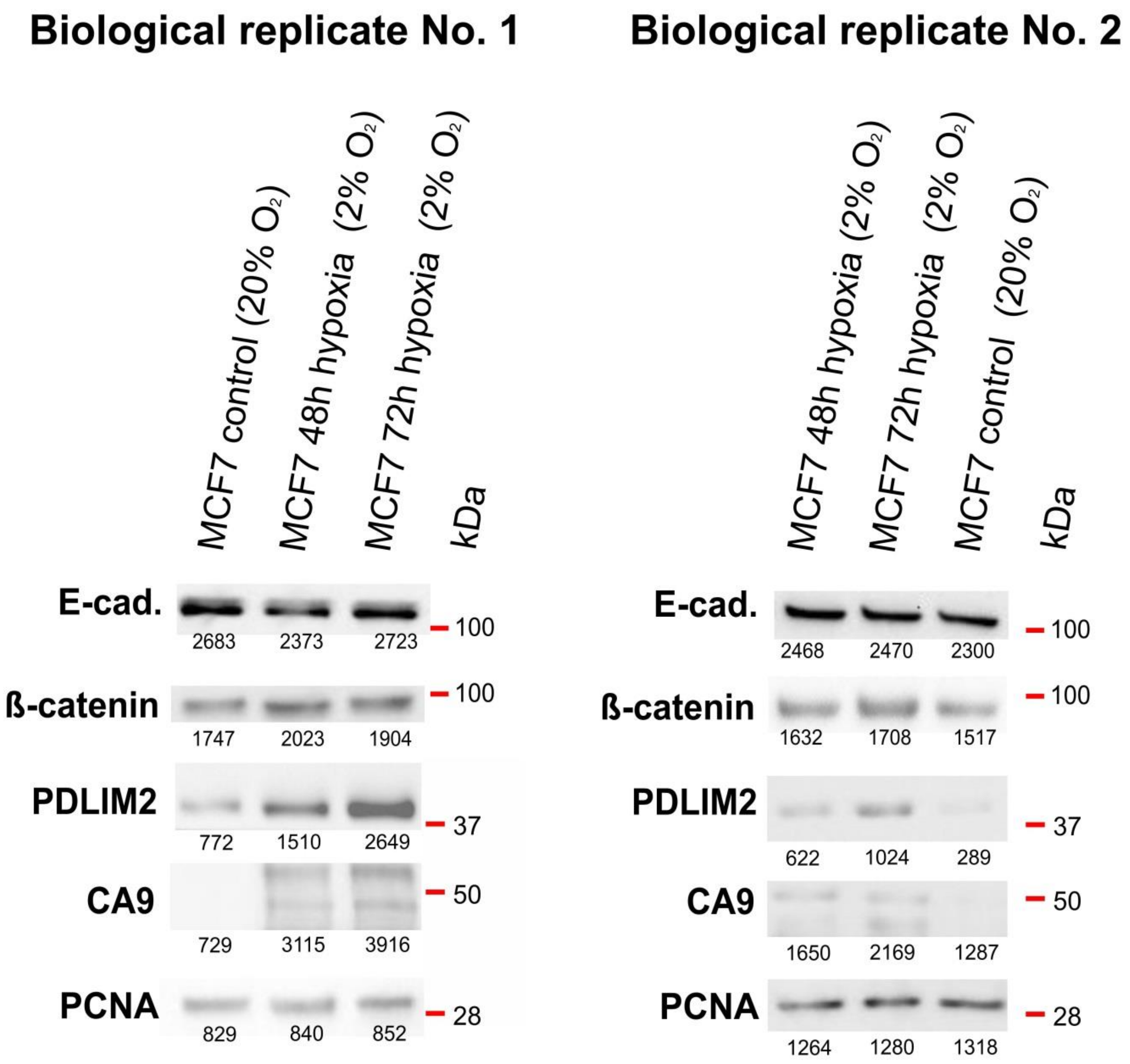


**Figure S5: Effects of short-term exposure to hypoxia (2% $O_2$ for 48 and 72 hours) on EMT markers and PDLIM2 protein levels in MCF7 cells. Related to Fig. 2B.** Carbonic anhydrase-9 (CA9) was monitored as a control marker for the induction of hypoxia. Short-term hypoxia led to increased levels of PDLIM2, and increased CA9 levels correlated with the duration of hypoxia. However, no changes of E-cad. and β-catenin protein levels indicating EMT induction was observed. PCNA was used as a loading control. Blots represent two independent experiments (biological replicates). Numbers under the protein bands represent their integral optical density (INT*$mm^2$).
