## Additional file 6 for "PDZ and LIM domain protein 2 plays dual and context-dependent roles in breast cancer development"

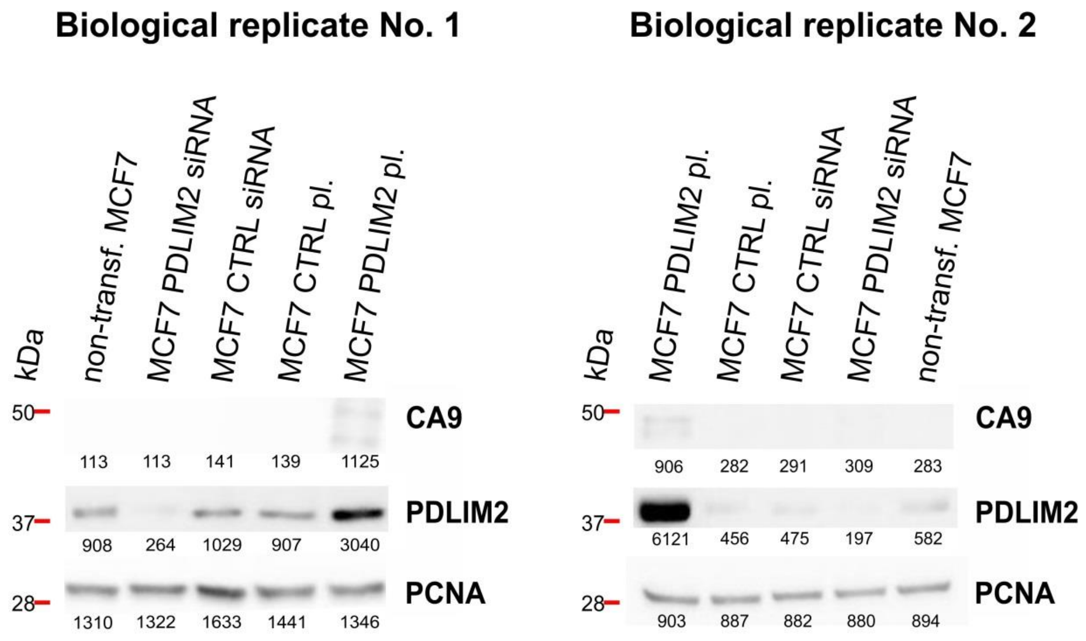


**Figure S6: Effects of altered PDLIM2 protein levels on the hypoxic marker carbonic anhydrase 9 (CA9) in MCF7 cells. Related to Fig. 2C.** PDLIM2 protein levels were altered by siRNA silencing (MCF7 PDLIM2 siRNA) compared to the control (MCF7 CTRL siRNA) or by PDLIM2 overexpression (MCF7 PDLIM2 pl.) compared to the control (MCF7 CTRL pl.) Overexpression of PDLIM2 led to increased CA9 levels. Blots represent two independent experiments (biological replicates). Numbers under the protein bands represent their integral optical density (INT*$mm^2$).
