## Additional file 7 for "PDZ and LIM domain protein 2 plays dual and context-dependent roles in breast cancer development"

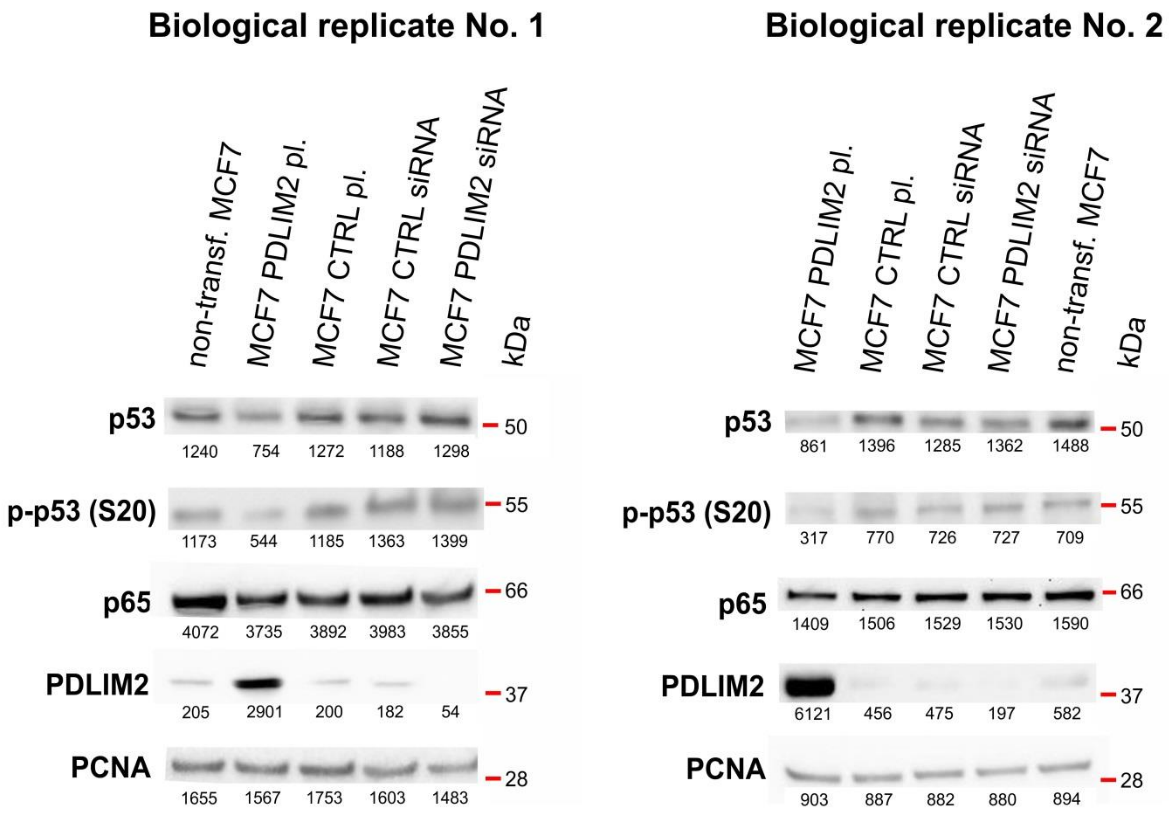


**Figure S7: Effects of altered PDLIM2 protein levels on p53 levels, p53 (S20) phosphorylation and p65 levels in MCF7 cells. Related to Fig. 2D.** PDLIM2 protein levels were altered by siRNA silencing (MCF7 PDLIM2 siRNA) compared to the control (MCF7 CTRL siRNA) or by PDLIM2 overexpression (MCF7 PDLIM2 pl.) compared to the control (MCF7 CTRL pl.). PDLIM2 overexpression led to decreased p53 levels and decreased p53 (S20) phosphorylation; PDLIM2 silencing had the opposite effect. No effect of PDLIM2 alterations on p65 protein levels was observed. Blots represent two independent experiments (biological replicates). Numbers under the protein bands represent their integral optical density (INT*mm$^2$).
