## Additional file 8 for "PDZ and LIM domain protein 2 plays dual and context-dependent roles in breast cancer development"

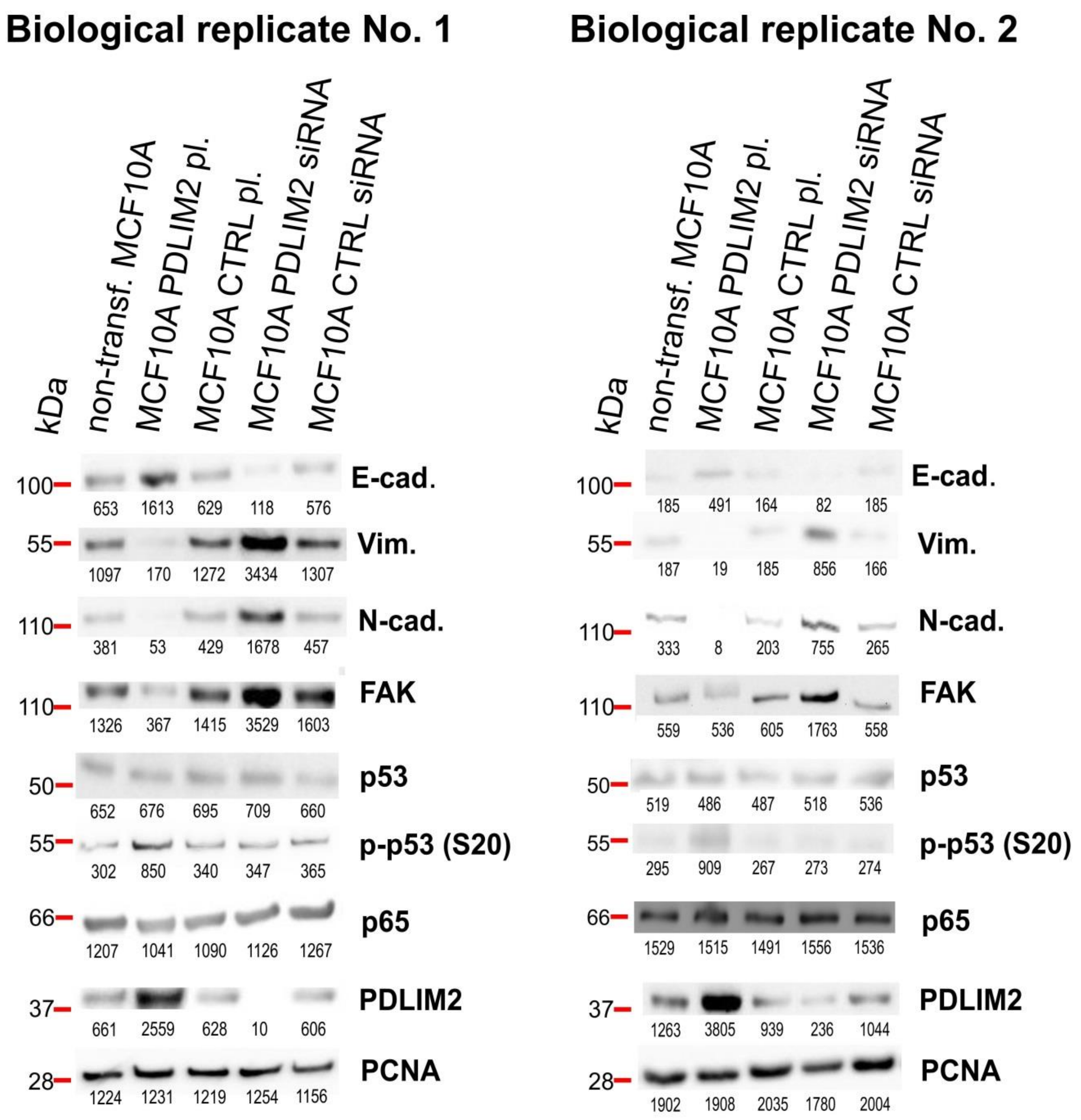


**Figure S8: Effect of altered PDLIM2 protein levels on EMT markers, p53, p-p53 (S20) and p65 in MCF10A cells. Related to Fig. 4A.** PDLIM2 protein levels were altered by siRNA silencing (MCF7 PDLIM2 siRNA) compared the control (MCF7 CTRL siRNA) or by PDLIM2 overexpression (MCF7 PDLIM2 pl.) compared to the control (MCF7 CTRL pl.). The EMT markers E-cadherin (E-cad.), vimentin (Vim.) and N-cadherin (N-cad.) as well as focal adhesion kinase (FAK), p53, the active form of p53 (p-p53 (S20)), and p65 were monitored. PDLIM2 overexpression led to increased levels of the epithelial marker E-cadherin and p53 (S20) phosphorylation and to decreased levels of the mesenchymal markers vimentin, N-cadherin and FAK. PDLIM2 silencing had the opposite effect. No effect of PDLIM2 alterations on p53 and p65 protein levels was observed. Blots represent two independent experiments (biological replicates). Numbers under the protein bands represent their integral optical density (INT*mm$^2$).
