## Additional file 9 for "PDZ and LIM domain protein 2 plays dual and context-dependent roles in breast cancer development"

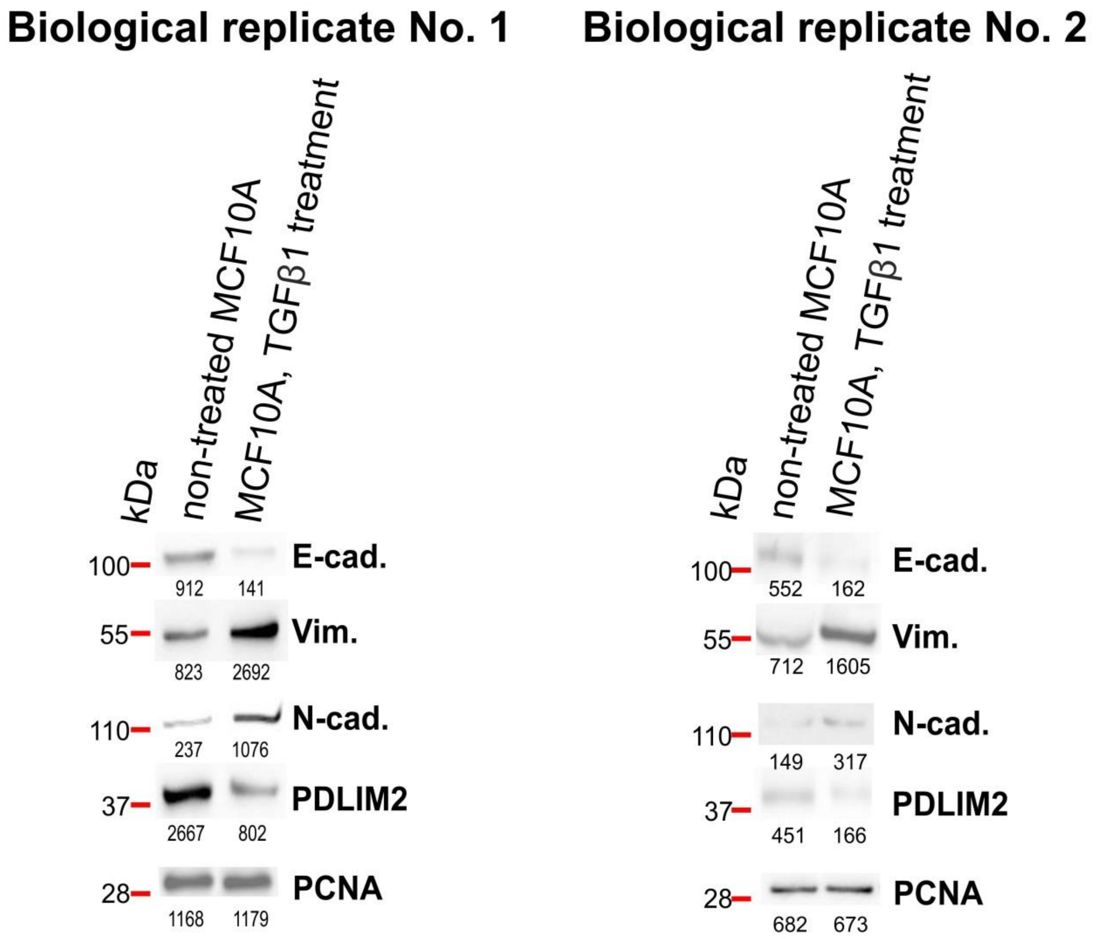


**Figure S9: Effects of EMT induction on PDLIM2 protein levels in MCF10A cells. Related to Fig. 4B.** EMT was induced by TGFβ1 treatment (1 ng/ml for 24 hours). The EMT markers E-cadherin (E-cad.), vimentin (Vim.) and N-cadherin (N-cad.) as well as focal adhesion kinase (FAK) were monitored. EMT induction led to decreased PDLIM2 levels, and successful EMT induction was confirmed by decreased levels of epithelial marker E-cadherin and increased levels of N-cadherin and vimentin. Blots represent two independent experiments (biological replicates). Numbers under the protein bands represent their integral optical density (INT*$mm^2$).
