## Additional file 10 for "PDZ and LIM domain protein 2 plays dual and context-dependent roles in breast cancer development"

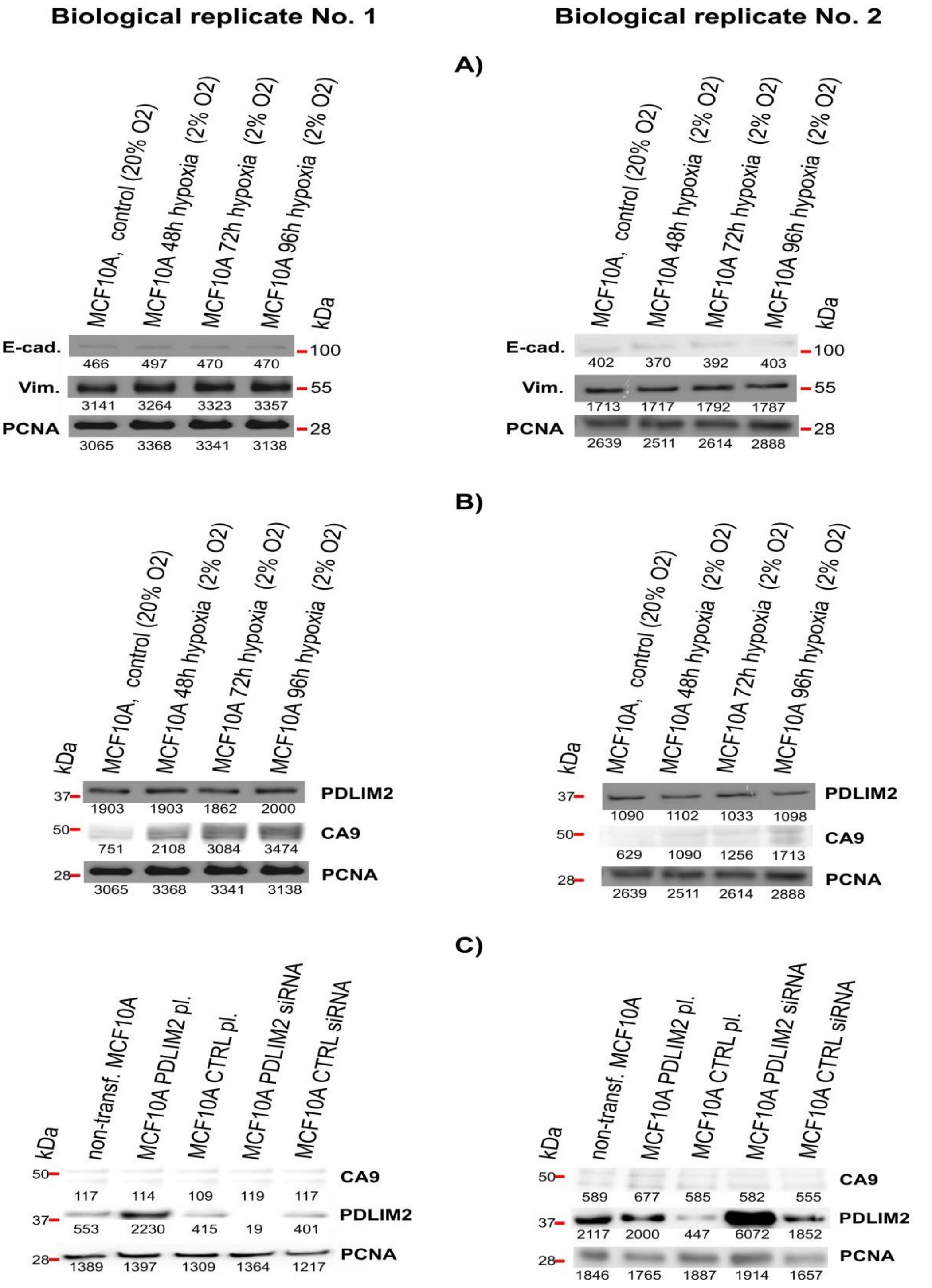


**Figure S10: Effect of short term and long term expositions to hypoxic conditions on EMT induction and effect of PDLIM2 alterations on CA9 in MCF10A cells. (A)** Neither short term nor long term expositions to hypoxic conditions (48, 72 and 96 hours) were able induce EMT in MCF10A cells (no changes in E-cad. and Vim. protein levels). **(B)** Hypoxic conditions have no effect on PDLIM2 protein levels, nonetheless, increased CA9 levels correlated with the duration of hypoxia. **(C)** PDLIM2 alterations have no effect on CA9 protein levels. Blots represent two independent experiments (biological replicates). Numbers under the protein bands represent their integral optical density (INT*mm$^2$).
