## Additional file 11 for "PDZ and LIM domain protein 2 plays dual and context-dependent roles in breast cancer development"

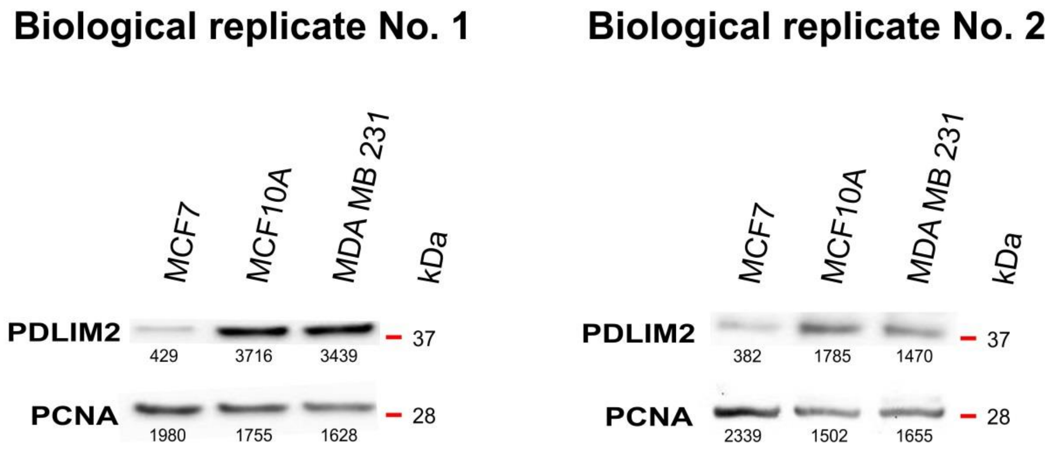


**Figure S11: PDLIM2 protein levels in different breast cell lines (MCF7, MCF10A and MDA MB 231).** The low-invasion MCF7 breast cancer cell line exhibited lower protein levels than both the immortalized normal epithelial breast cell line MCF10A and the highly invasive triple-negative breast cancer cell line MDA MB 231. Blots represent two independent experiments (biological replicates). Numbers under the protein bands represent their integral optical density (INT*mm$^2$).
