## Additional file 12 for "PDZ and LIM domain protein 2 plays dual and context-dependent roles in breast cancer development"

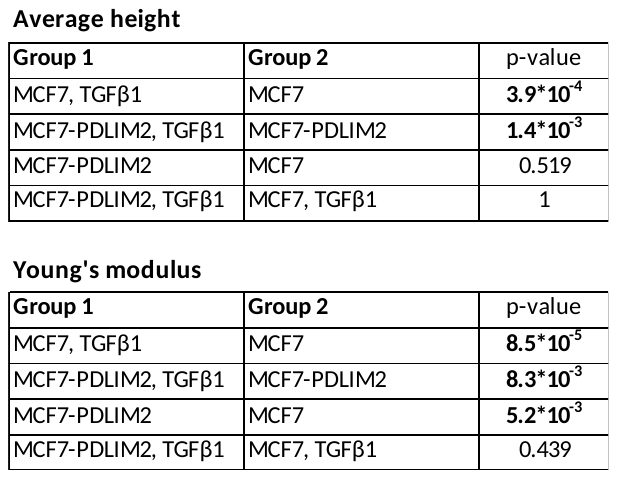

**Table S1: Statistics of average height and Young’s modulus of the cells measured by AFM**. **Related to Fig. 1E.** MCF7 and MCF7-PDLIM2 cells after or without TGFβ1 treatment, all comparisons. Eleven AFM measurements per group. Significant changes are highlighted in **bold**. See Fig. 1E for boxplot visualization.
