## Additional file 13 for "PDZ and LIM domain protein 2 plays dual and context-dependent roles in breast cancer development"

|  | protein | | transcript | |
| --- | --- | --- | --- | --- |
|  | fold change | p-value | fold change | p-value |
| G1: N1-2/N0 | **1.30** | **0.009** | **1.287** | **0.007** |
| G3:N1-2/N0 | 1.06 | 0.866 | 0.851 | 0.215 |
| N1-2/N0 | 1.18 | 0.415 | 1.047 | 0.571 |
| G3/G1 | **0.71** | **0.008** | 1.038 | 0.645 |
| N1-2:G3/G1 | 0.64 | 0.063 | 0.844 | 0.109 |
| ER+/ER- | 1.42 | 0.208 | 1.159 | 0.258 |
| HER2+/HER2- | 0.65 | 0.060 | 1.056 | 0.671 |

**Table S2: Connection between PDLIM2 and** **clinicopathological parameters of breast cancer based on our previous combined proteomics and transcriptomics study (n=96 in total) [5].** The data show that higher protein and transcript levels of PDLIM2 are connected with (i) lymph node metastasis of luminal A grade 1 tumors and (ii) with low tumor grade. Significant changes in protein levels and expression (according to the criteria defined in the Experimental Procedures section of [5]) are indicated in **bold**.
