## Additional file 14 for "PDZ and LIM domain protein 2 plays dual and context-dependent roles in breast cancer development"

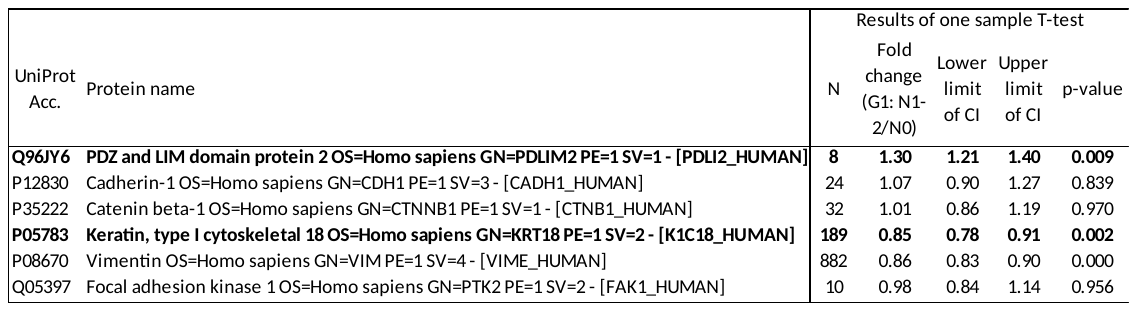

**Table S3: iTRAQ-2DLC-MS/MS quantitative protein-level data for PDLIM2 and epithelial and mesenchymal marker proteins in lymph node-positive (n=24) vs. negative (n=24) luminal A grade 1 tumors in our previous combined proteomics and transcriptomics study (n=96 in total) [5].** The data show that PDLIM2 was upregulated, while KRT18 was downregulated in lymph node-positive vs. negative tumor**s**. N, number of peptide-level observations; CI, confidence interval. Significant changes in protein levels and expression (according to criteria defined in the Experimental Procedures section of [5]) are indicated in **bold**.
