## Additional file 15 for "PDZ and LIM domain protein 2 plays dual and context-dependent roles in breast cancer development"

**Methods S1: Young's modulus mapping by Atomic Force Microscopy**

A standard bio AFM microscope JPK NanoWizard 3 from JPK (JPK, Berlin, Germany) was used to perform the force mapping procedure. The scanning-by-probe head (maximal visualization range 100-100-15 µm in X-Y-Z axis) of the AFM microscope was placed on an inverted optical microscope Olympus IX-81, and a 10x objective was used to find the area covered with cells and to place the cantilever in the proper position for the force mapping procedure. A plastic Petri dish i.d. 34 mm (TPP, Trasadingen, Switzerland) either with distilled water for instrument calibration or with cultured cells was placed inside the Petri dish heater (JPK) preheated to 37 °C.

A non-coated silicon nitride AFM probe Hydra 2R-100N (AppNano, Mountain View, CA, United States) equipped with a pyramidal silicon tip was used for all experiments. The probe was calibrated prior to every set of experiments as described below.

The calibration procedure was done in water, when the whole setup was preheated (Petri dish heater, JPK) to 37 °C for 30 min. Afterwards, the laser reflection sum was maximized, followed by centering of the laser detector. The AFM probe was introduced to contact with the surface during a standard process of landing. The sensitivity of the AFM setup was determined as a slope of the force-distance curve measured by lifting the cantilever with a Z-height of 450 nm, and the acquisition time per curve was 2 s. The sensitivity was in the range of 13.37-16.09 nm/V, and cantilever stiffness was calibrated by measurement of its thermal noise and proceeded between 18.24 and 20.20 nN/m on different days of experiments.

The bio AFM setting was identical for all force mapping procedures. The setpoint value was 2.5 nN (relative to baseline value), time per curve 0.5 s, Z-length 2.0 µm, speed of curve recording 4.0 µm/s, and the force-distance curves were recorded with a data sample rate of 2 kHz. The force mapping procedure was performed as a step-by-step recording of force-distance curves in the network of 64x64 points on a 100x100 $\mu m^2$ area. The whole content of the Petri dish, i.e., 2 ml of medium, was exchanged for fresh medium (preheated and $CO_2$ equilibrated in a standard $CO_2$ incubator) using a sterile plastic syringe connected to the inlet tube of the Petri dish holder (JPK).

The force mapping process provides a network of force-distance curves (FDC, dependency of tip-sample interaction force on tip height above the surface), so-called force maps (FM). The absolute value of Young's modulus can be determined by fitting the FDC by the Bilodeau model [18].

For the processing of FMs, AtomicJ software was used, and a robust exhaustive contact estimator and classical model fit L2 were employed to fit the approach curve. A Poisson ratio of 0.5 (incompressible material) and 2$^{nd}$ baseline degree were used as fitting parameters. Final visualization of the images and stiffness maps was performed with Gwyddion software ver. 2.44 [20].
