## Additional file 16 for "PDZ and LIM domain protein 2 plays dual and context-dependent roles in breast cancer development"

**Dataset S1: AFM data for all measurements –**

11 biological replicates per group

A. untreated MCF7 cells - cell height

B. TGFβ1 treated MCF7 cells - cell height

C. untreated MCF7-PDLIM2 cells - cell height

D. TGFβ1 treated MCF7-PDLIM2 cells - cell height

E. untreated MCF7 cells - Young's modulus

F. TGFβ1 treated MCF7 cells - Young's modulus

G. untreated MCF7-PDLIM2 cells - Young's modulus

H. TGFβ1 treated MCF7-PDLIM2 cells - Young's modulus

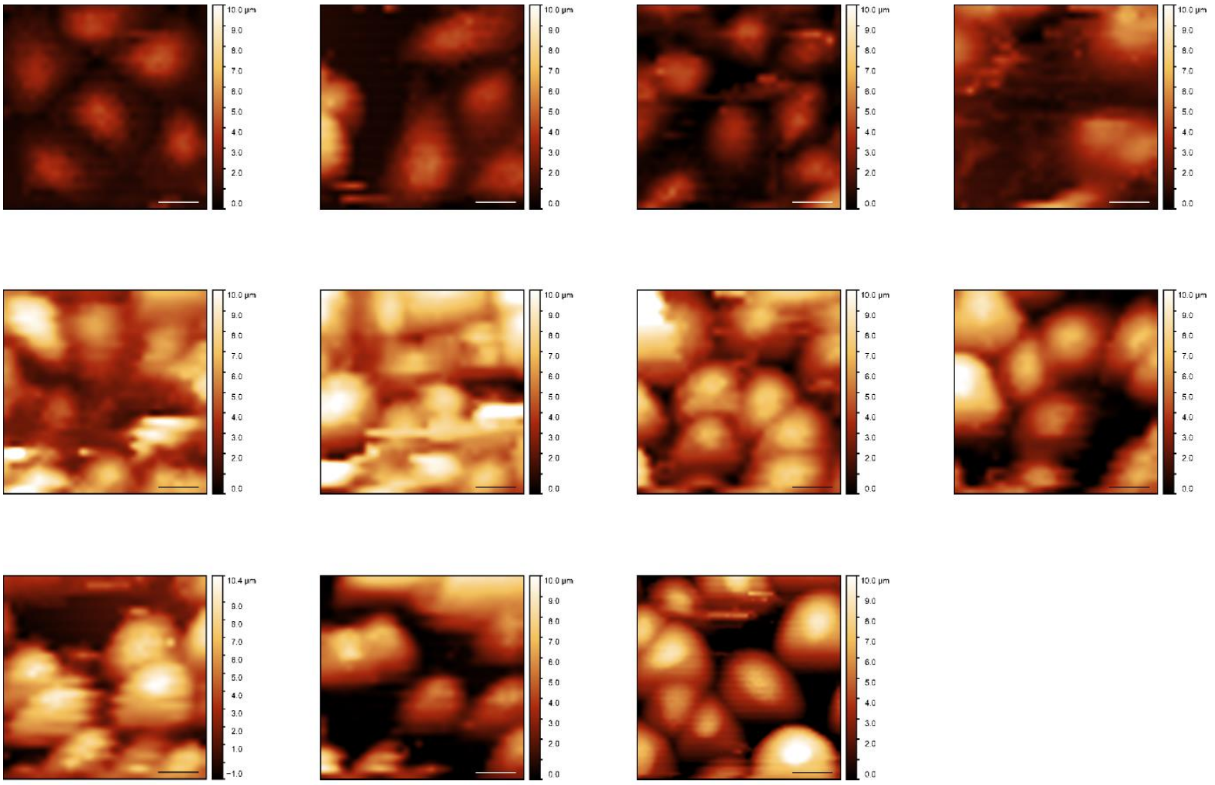

A. Distribution of untreated MCF7 cells height over the cells.
Scale bar inside the image is equal to 20 µm.

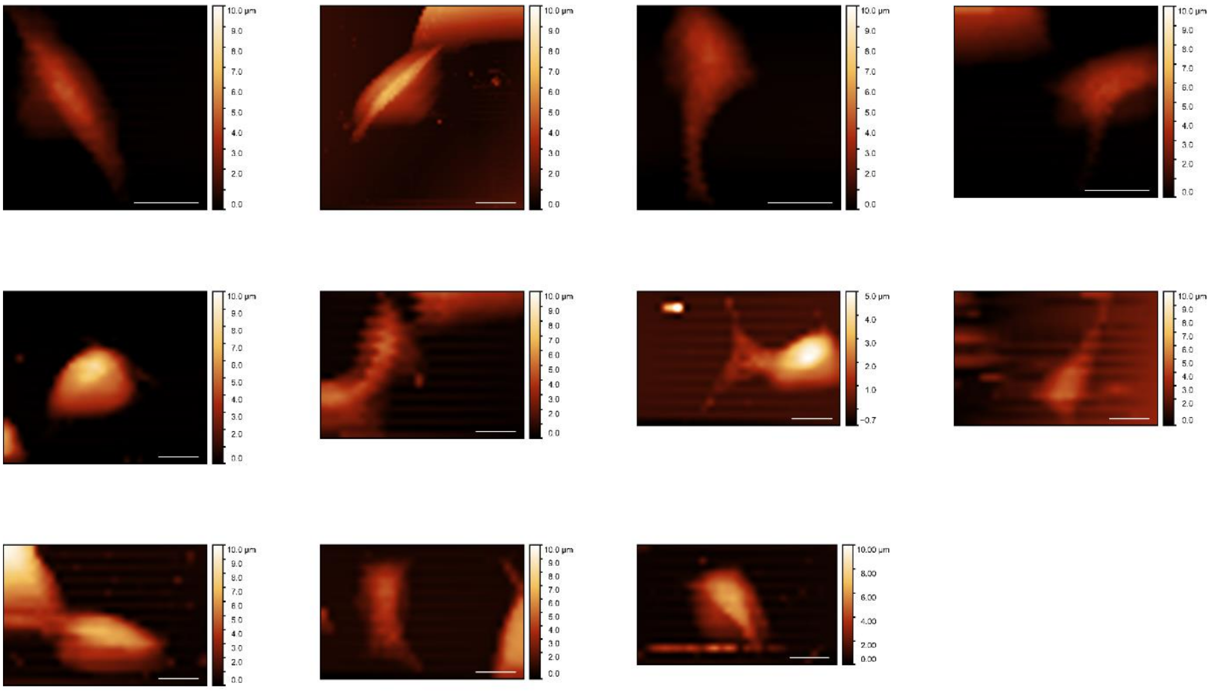

B. Distribution of TGFβ1 treated MCF7 cells height over the cells.
Scale bar inside the image is equal to 20 µm.

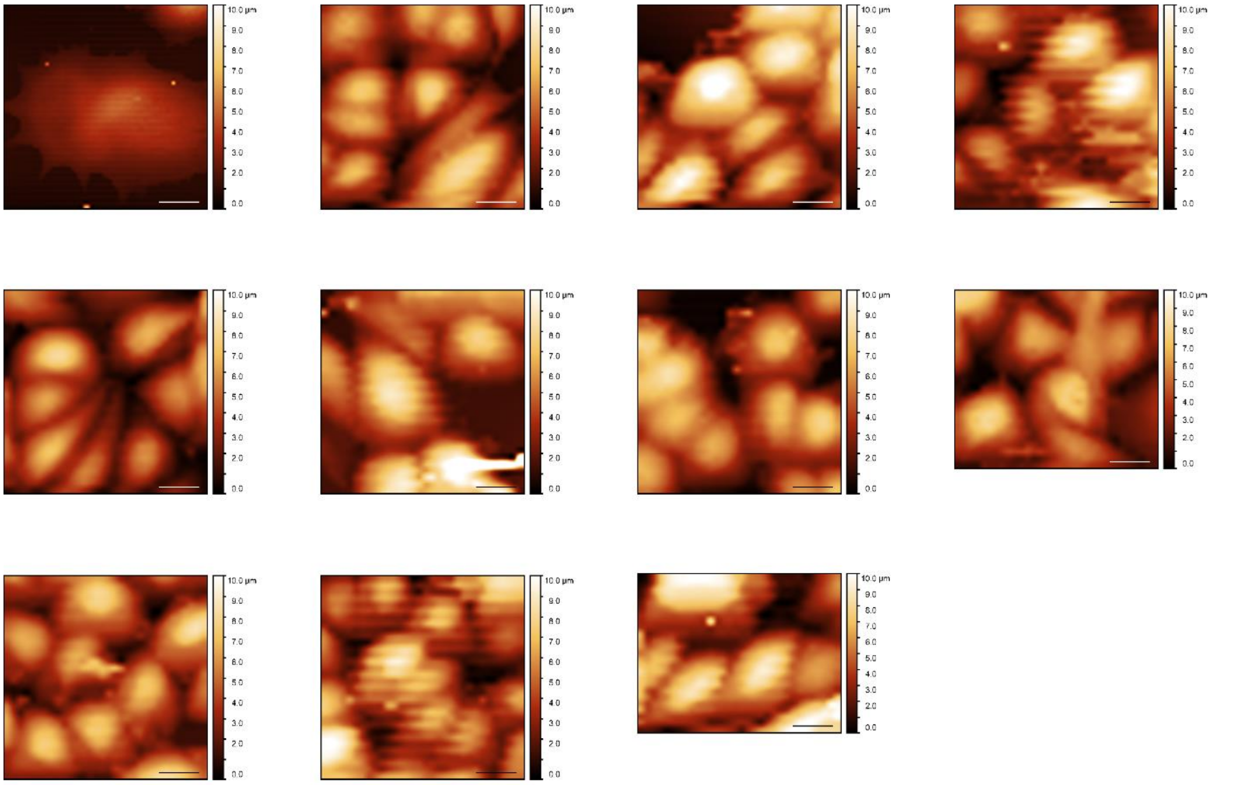

C. Distribution of untreated MCF7-PDLIM2 cells height over the cells.
scale bar inside the image is equal to 20 µm.

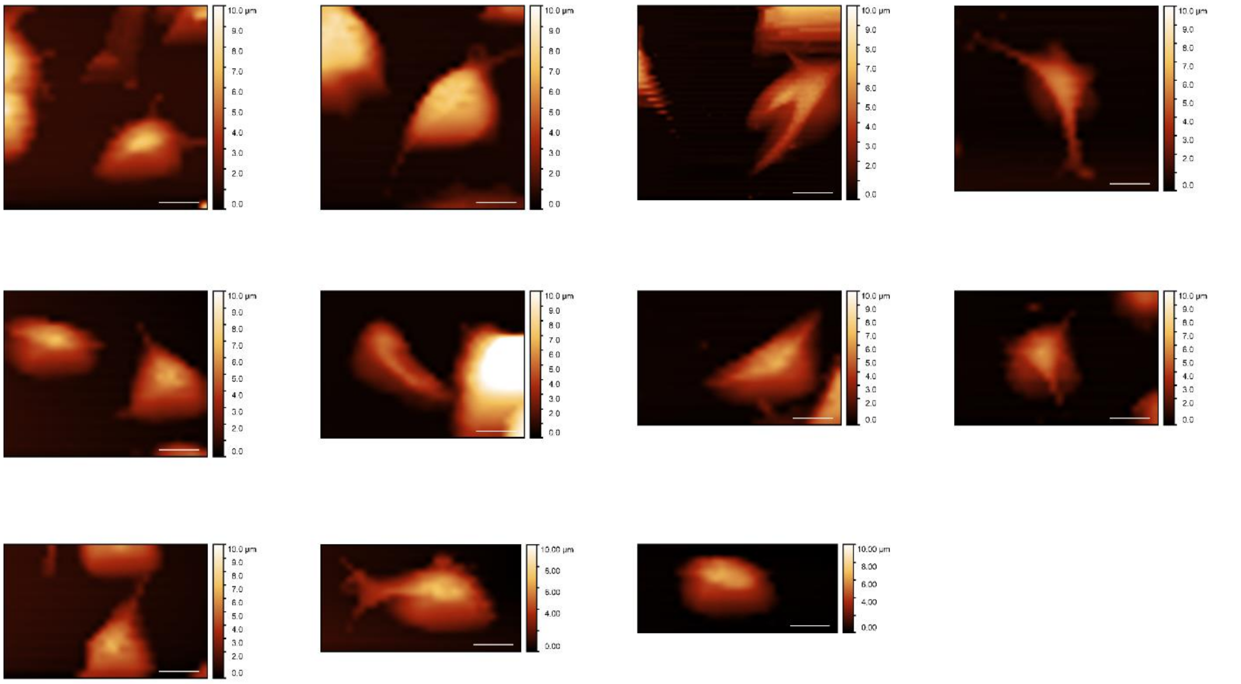

D. Distribution of TGFβ1 treated MCF7-PDLIM2 cells height over the cells.
Scale bar inside the image is equal to 20 µm.

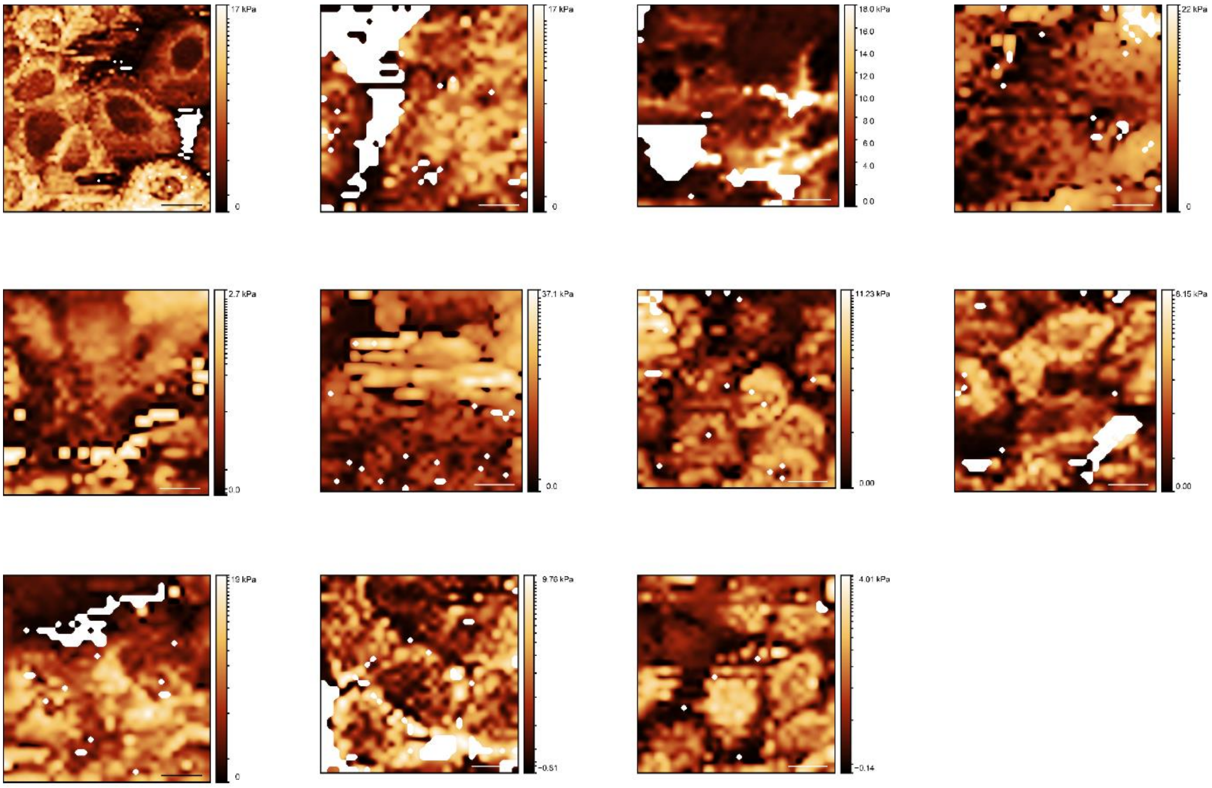

E. Distribution of untreated MCF7 cells Young's modulus over the cells.
Scale bar inside the image is equal to 20 μm.

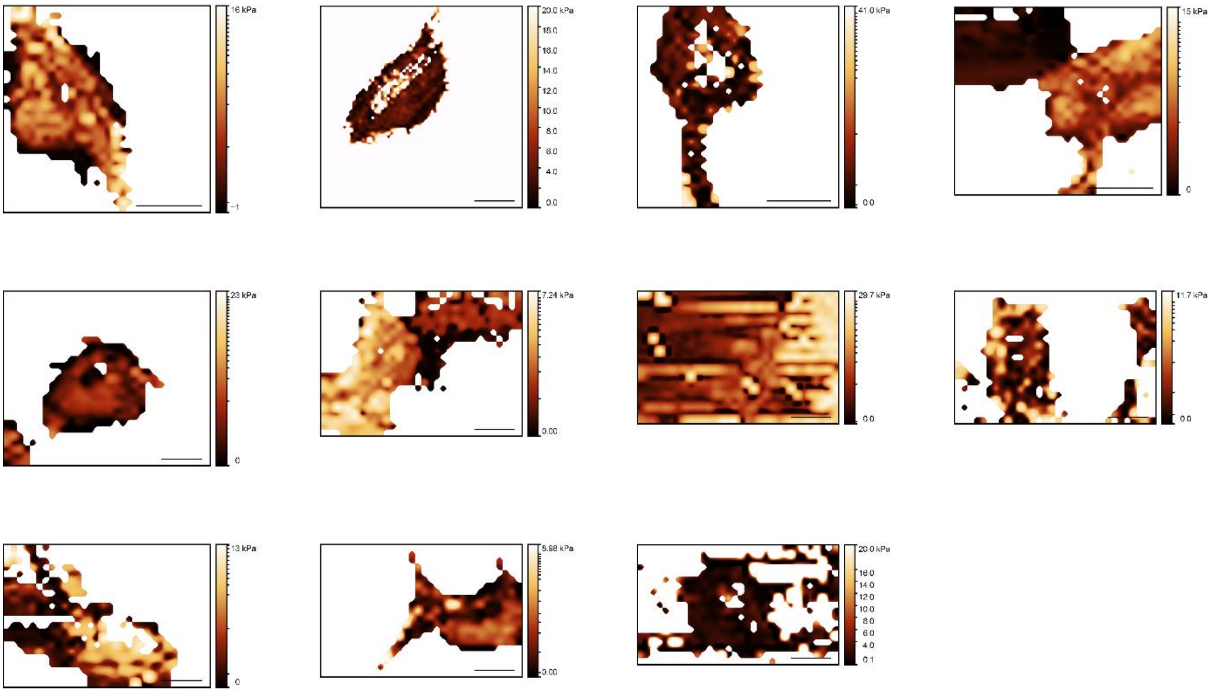

F. Distribution of TGFβ1 treated MCF7 cells Young's modulus over the cells.
Scale bar inside the image is equal to 20 μm.

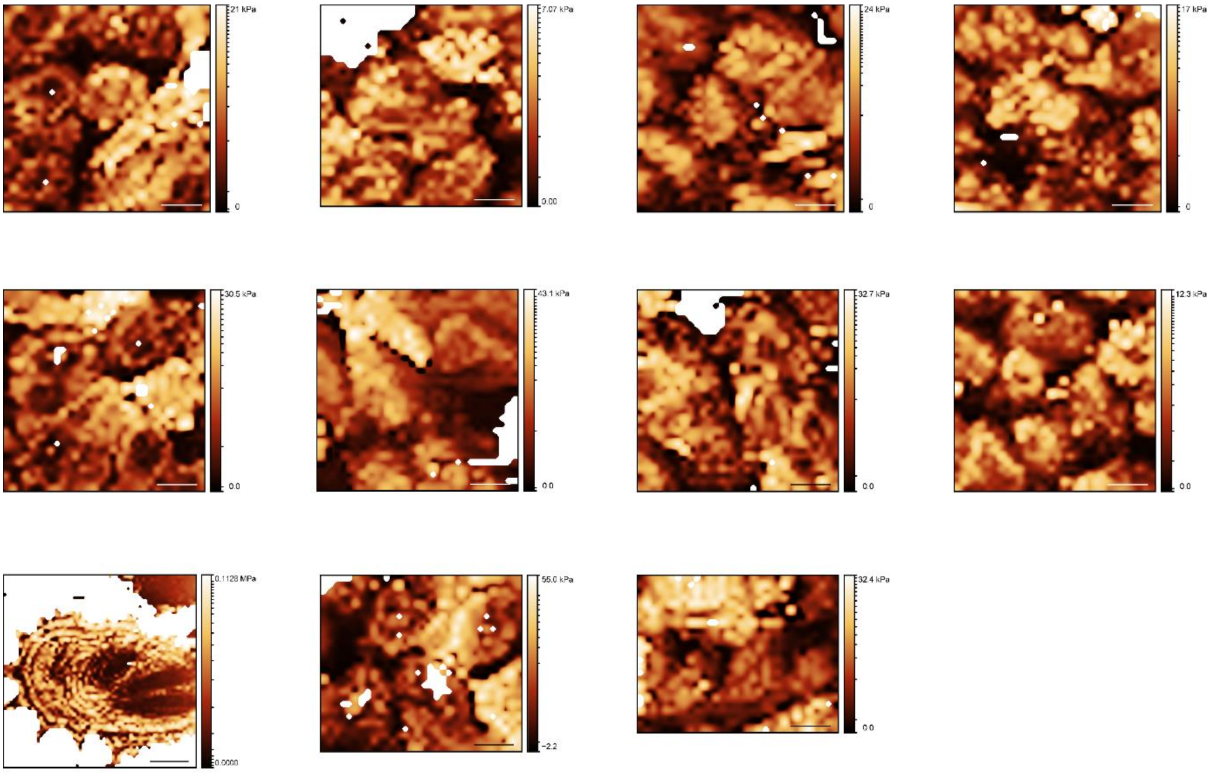

G. Distribution of untreated MCF7-PDLIM2 cells Young's modulus over the cells.
Scale bar inside the image is equal to 20 μm.

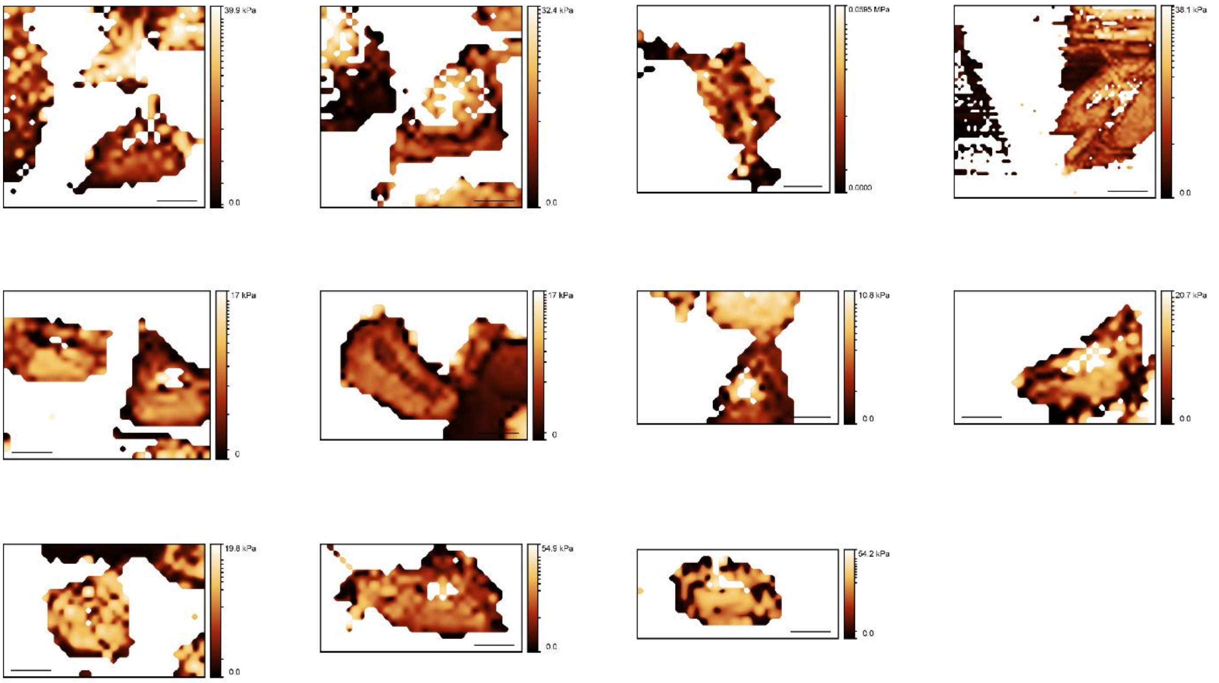

H. Distribution of TGFβ1 treated MCF7-PDLIM2 cells Young's modulus over the cells.
Scale bar inside the image is equal to 20 μm.
