## Additional file 17 for "PDZ and LIM domain protein 2 plays dual and context-dependent roles in breast cancer development"

**Dataset S2: xCELLigence data for MCF7 migration and invasion measurements**
- MCF7 cells with altered PDLIM2 protein levels (two biological replicates)

**I.)** Biological replicate No. 1 (see Fig. 3A and 3B in the manuscript)
**II.)** Biological replicate No. 2

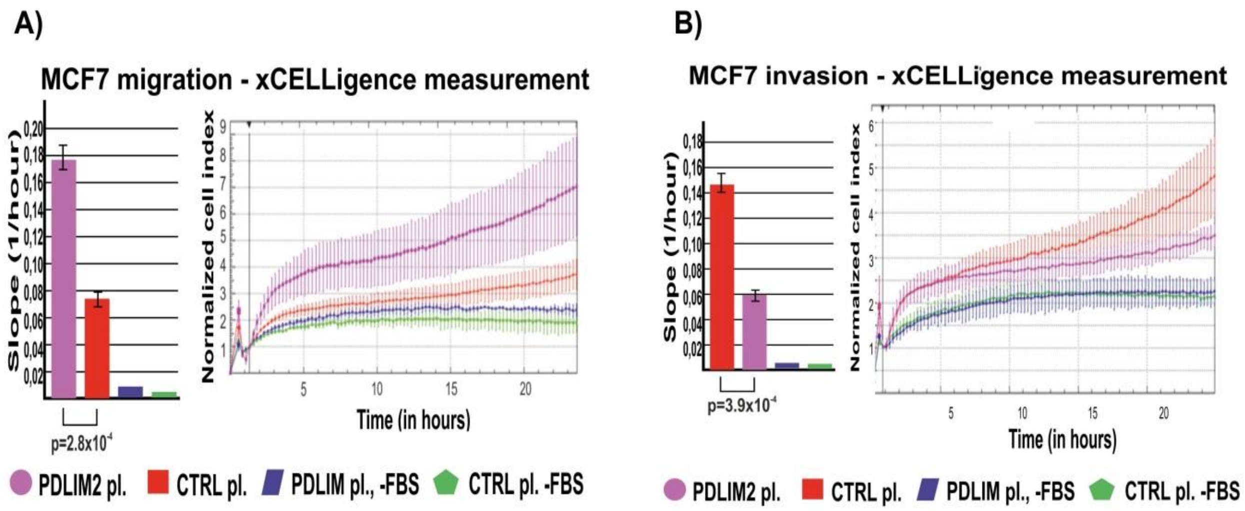


**I.) (A)** Effect of PDLIM2 overexpression (PDLIM2 pl.) on migration of MCF7 cells (in comparison with control cells with endogenous PDLIM2 levels (CTRL pl.)) measured by xCELLigence system. **(B)** Effect of PDLIM2 overexpression (PDLIM2 pl.) on invasiveness of MCF7 cells (in comparison with control cells with endogenous PDLIM2 levels (CTRL pl.)).

A)

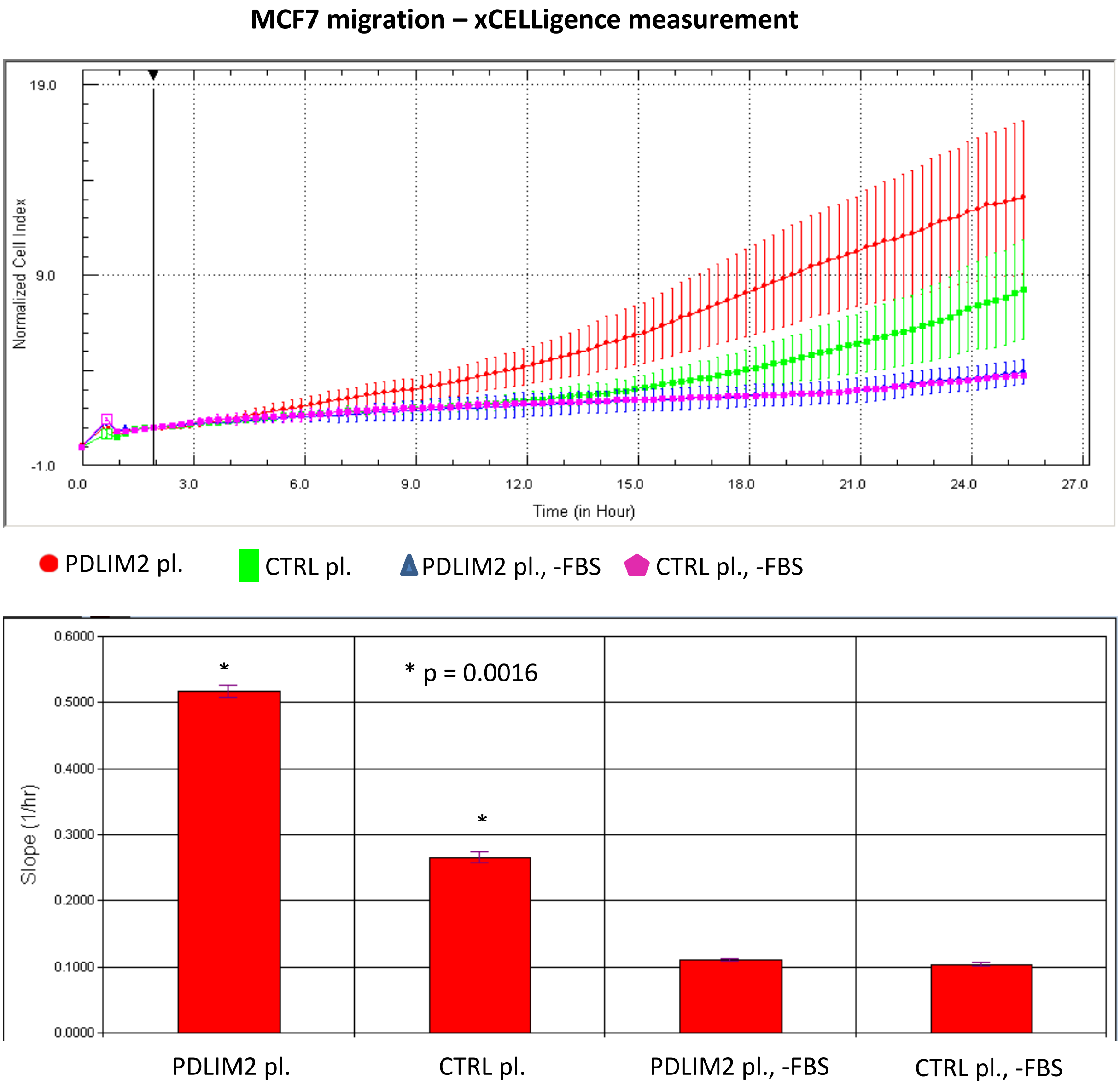

MCF7 migration – xCELLigence measurement
19.0
9.0
-1.0
Normalized Cell Index
0.0
3.0
6.0
9.0
12.0
15.0
18.0
21.0
24.0
27.0
Time (in Hour)
PDLIM2 pl.
CTRL pl.
PDLIM2 pl., -FBS
CTRL pl., -FBS
0.6000
0.5000
0.4000
0.3000
0.2000
0.1000
0.0000
Slope (1/hr)
*
* p = 0.0016
PDLIM2 pl.
CTRL pl.
PDLIM2 pl., -FBS
CTRL pl., -FBS

**B)**

**MCF7 invasion – xCELLigence measurement**

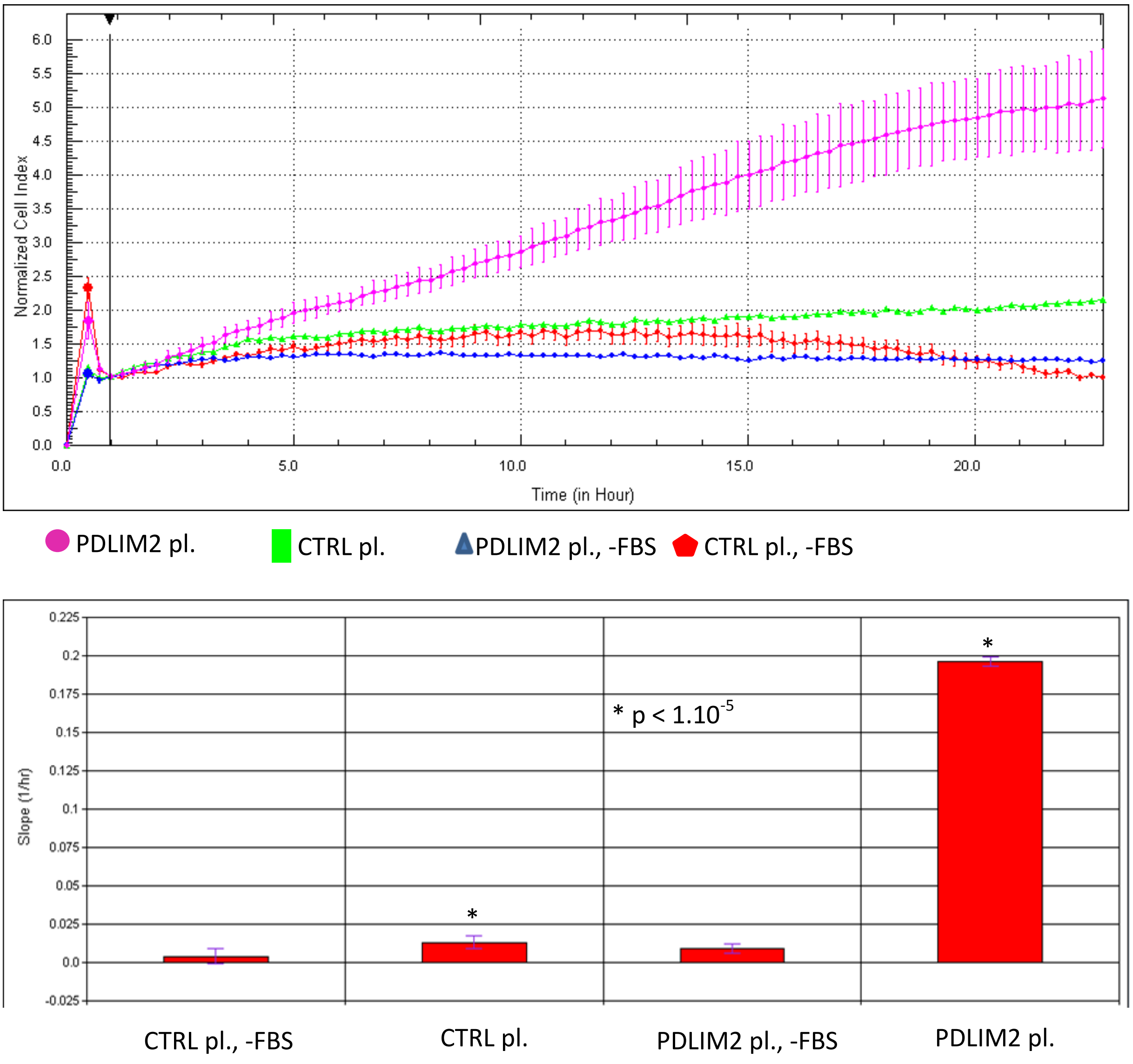


**<u>II.)</u> (A)** Effect of PDLIM2 overexpression (PDLIM2 pl.) on migration of MCF7 cells (in comparison with control cells with endogenous PDLIM2 levels (CTRL pl.)) measured by xCELLigence system. **(B)** Effect of PDLIM2 overexpression (PDLIM2 pl.) on invasiveness of MCF7 cells (in comparison with control cells with endogenous PDLIM2 levels (CTRL pl.)).
